# Structural and functional determinants inferred from deep mutational scans

**DOI:** 10.1101/2022.02.21.481196

**Authors:** Priyanka Bajaj, Kavyashree Manjunath, Raghavan Varadarajan

## Abstract

Mutations that affect protein binding to a cognate partner primarily occur either at buried residues or at exposed residues directly involved in partner binding. Distinguishing between these two categories based solely on mutational phenotypes is challenging. The bacterial toxin CcdB kills cells by binding to DNA Gyrase. Cell death is prevented by binding to its cognate antitoxin CcdA, at an extended interface that partially overlaps with the GyrA binding site. Using the CcdAB toxin-antitoxin (TA) system as a model, a comprehensive site-saturation mutagenesis library of CcdB was generated in its native operonic context. The mutational sensitivity of each mutant was estimated by evaluating the relative abundance of each mutant in two strains, one resistant and the other sensitive to the toxic activity of the CcdB toxin, through deep sequencing. The ability to bind CcdA was inferred through a RelE reporter gene assay, since the CcdAB complex binds to its own promoter, repressing transcription. By analysing mutant phenotypes in the CcdB sensitive, CcdB resistant and RelE reporter strains, it was possible to assign residues to buried, CcdA interacting or GyrA interacting sites. A few mutants were individually constructed, expressed, and biophysically characterised to validate molecular mechanisms responsible for the observed phenotypes. Residues inferred to be important for antitoxin binding, are also likely to be important for rejuvenating CcdB from the CcdB-Gyrase complex. Therefore, even in the absence of structural information, when coupled to appropriate genetic screens, such high-throughput strategies can be deployed for predicting structural and functional determinants of proteins.

**Broader Impact Statement:** Partial loss-of-function mutations predominantly occur either at buried-site or exposed, active-site residues. We report a facile method to identify multiple binding sites for different interacting partners for a protein, and distinguish them from buried site and exposed non active-site residues, solely from mutational data.

## INTRODUCTION

The amino acid sequence dictates the tertiary structure of a protein which is closely tied to its activity. From a wealth of previous studies, it is known that mutations that affect function primarily occur either at exposed active site/ligand binding residues or at buried sites important for protein stability^1, 2^. However, distinguishing them, purely from mutational phenotypes is challenging. Several computational approaches exploit sequence-structure relationship to predict functional patches on the protein surface through *in silico* modelling based on a query protein sequence, sequence conservation^3^ or structural homology with well-characterised proteins^4^. This is challenging for proteins with low sequence identity to those present in sequence databases^5^. Traditional methods are low-throughput and require purification and characterisation of large numbers of individual variants to identify active site residues such as protein: protein and protein: ligand binding site residues^6^. X-ray crystallography, NMR spectroscopy, and cryo-electron microscopy of a protein complex can be used to identify residues important in binding to its interacting partner, a prerequisite is high yield and homogenous preparation of purified protein^7–9^. While these methods are useful in providing atomic level information but are labour intensive and not easily parallelizable. The advent of next generation sequencing has revolutionised biology^1, 10^, resulting in the development of various approaches that can be used to predict structural features in the absence of a structure^2, 11^. Alanine scanning, and cysteine scanning mutagenesis approaches have been previously used to predict functional residues but have limitations. Alanine scanning mutagenesis is laborious as each alanine substituted protein needs to be individually expressed and characterised^12^. Cysteine scanning mutagenesis requires an additional step of labelling the exposed residues, effects and labelling of a buried cysteine may result in misfolding of the protein, thereby resulting in production of false positive results^13^. Generally, buried residues as well as active site residues are sensitive to mutations but differ in their mutational tolerance to aliphatic and charged substitutions^2^. Deep mutational scanning is a promising tool for mapping sequence-activity relationships in proteins for which an observable phenotypic readout is available. Mutations at the active site generally affect the specific activity of the protein as the native conformation remains intact, while buried site mutants affect the stability and folding of the protein, thereby affecting the total activity of the protein. However, distinguishing between these two classes of residues solely from phenotypic data is challenging^14^. The situation is even more complex for proteins with multiple binding partners.

The system used in the present study is a *ccd* operon. This is a Type II toxin-antitoxin (TA) system and both *ccdA* and *ccdB* genes encode proteins. CcdA acts as an antidote which neutralises the toxicity of CcdB by forming a tight complex with it, and also rejuvenating Gyrase from its complex with CcdB^15^. The (CcdA)_2_-(CcdB)_2_ complex represses the operon at the transcriptional level by binding to the operator-promoter region of the operon to maintain the CcdB:CcdA ratio <1^16^. Under stress conditions, CcdA is degraded and the CcdB:CcdA ratio is increased. This causes transient de-repression of the operon and fresh synthesis of both CcdA and CcdB (Figure S1). The crystal structure, 3G7Z shows that the C-terminal intrinsically disordered region of CcdA binds consecutively to two overlapping sites of CcdB with different affinities. Both sites are important for rejuvenation and autoregulation of expression of the *ccd* operon^17, 18^. Promoter-operator binding of the CcdA-CcdB complex *in vivo* can be probed by co-expressing the complex and a reporter gene downstream of the *ccd* promoter within the cell. Dual selection reporter systems such as the tetA gene^19^, tetA-sacB cassette^20^, kill gene^21^ and RelE gene^22^ have previously been used for genome recombineering studies. The RelE reporter system is reported to achieve higher selection stringency than previously reported negative selection systems, usable in *E. coli* strains^22^.

CcdAB is a convenient system to study mutational effects in an operonic context. In this report we describe comprehensive single-site mutational scanning of CcdB in its operonic context, with the goal of determining multiple binding sites and, distinguishing active site residues from the buried site and exposed non active-site residues. We attempt to address the following issues: (1) Is identifying active site residues of CcdB solely from mutational phenotypes possible? (2) Can we distinguish Gyrase binding site residues from buried residues from mutational phenotypes? (3) How well does the amount of Accessible Surface Area buried upon complex formation with the interacting partner explain the mutational landscape of active site residues? (4) Is there any consistent pattern in substitution preferences at the active site residues? (5) Can we delineate molecular mechanisms behind the observed phenotypes in a high-throughput manner? (6) How can the inferred molecular mechanisms be validated? (7) Are the CcdA interacting residues impaired in CcdA binding also important for rejuvenating CcdB from the CcdB-Gyrase complex. Crystal structures of CcdB complexed to CcdA (PDBid: 3G7Z)^17^ and DNA Gyrase (PDBid: 1X75)^23^ were used to rationalise mutant phenotypes obtained from deep sequencing.

## RESULTS

### Deep sequencing of CcdB SSM library in operonic context

A comprehensive site-saturation mutagenesis (SSM) library of CcdB was prepared in its native operon that contained the promoter, *ccdA* and *ccdB* genes in pUC57 plasmid (a high copy number plasmid), to get an amplified response required to distinguish the mutant phenotype from WT. The pooled mutant library of CcdB, transformed in two strains, one resistant (resistant strain) and the other sensitive (sensitive strain) to the toxic activity of CcdB (Figure S2 A), was subjected to deep sequencing and each mutant analysed was assigned a variant score in the form of ‘Relative Fitness^CcdB^’ (RF^CcdB^) (Eq. 6, see methods). Out of ∼3200 (100 positions*32 codons) mutants expected by NNK codon mutagenesis of the *ccdB* gene, reads for ∼2700 mutants were available in the resistant strain (unselected library). Deep sequencing data from the two biological replicates were compared using different read cut-offs (Figure S2 B). The highest correlation of ∼0.96 between the two was obtained when the read cut-off in the resistant strain was taken as 100 (Figure 1 A). CcdB mutants were ranked based on their activity, for which the phenotypic readout is cell growth versus cell death. We first validated the deep sequencing data in a high-throughput manner by overlaying the RF^CcdB^ scores obtained by all CcdB mutants with synonymous CcdB mutants and non-functional CcdB mutants (Figure 1 B). The synonymous mutant dataset represents an internal positive control with the median value of ∼0.8 and the major fraction lie within a 2-fold range of the WT score (RF^CcdB^=1), i.e., between 0.5 to 2, suggesting these mutants show a phenotype similar to WT. In contrast, stop codon mutant dataset represents an internal negative control with median value of 16 and most non-sense mutations exhibited inactive phenotypes (RF^CcdB^>2).

**Figure 1:**
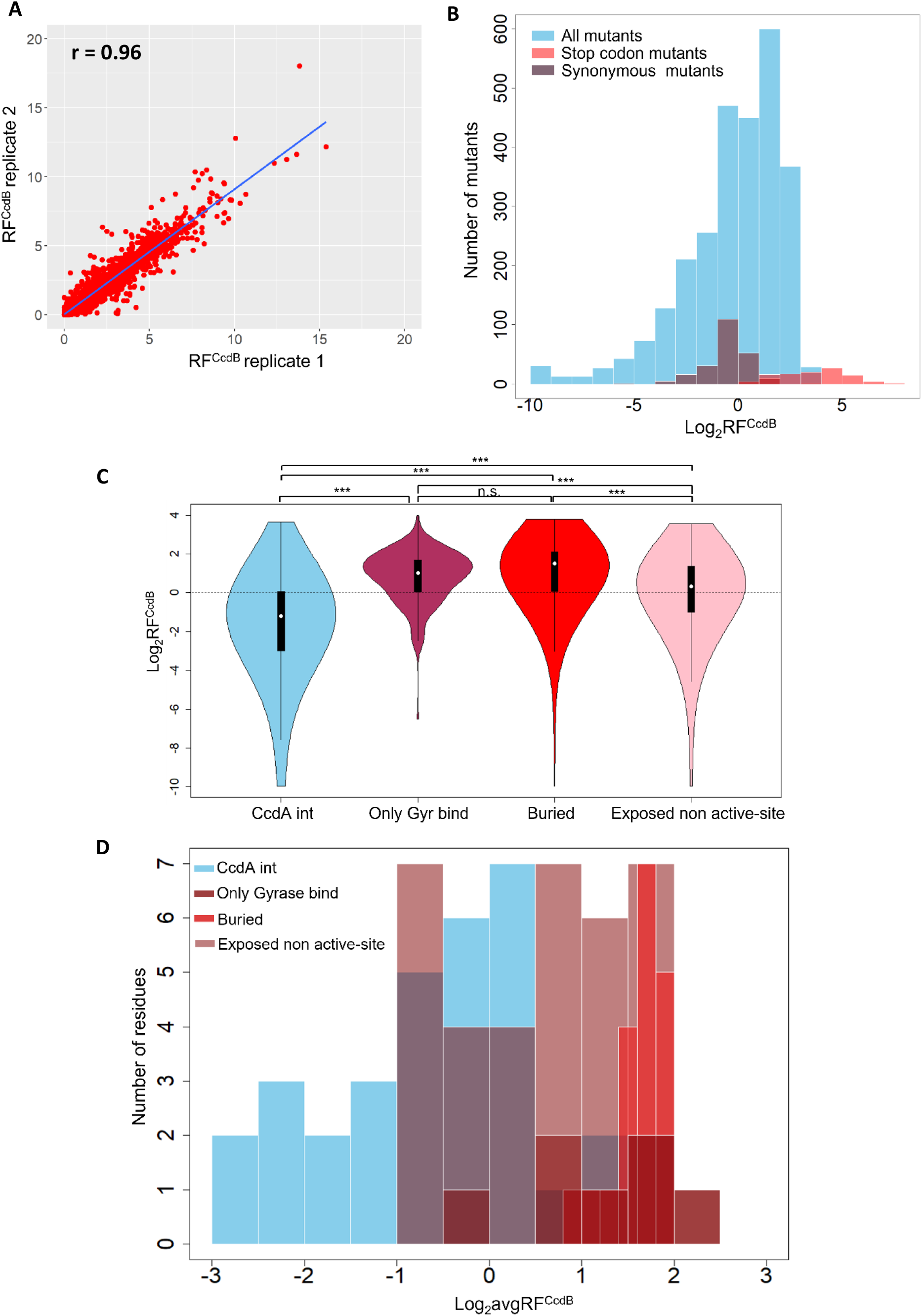
Reproducibility and distribution of Relative Fitness^CcdB^ (RF^CcdB^) values of the entire dataset. (A) Correlation between RF^CcdB^ values for the two biological replicates for mutants with read cut-off greater than 100 in the resistant strain Top10Gyr. (B) Histogram of RF^CcdB^ for all the mutants in the entire dataset (blue) with synonymous mutants (grey) and stop-codon mutants (pink). Overlapping region between the two classes is shown in light purple. (C) Violin plot with width proportional to the number of mutants at a given RF^CcdB^ value found in each class (mentioned on the x-axis). White dot in the middle of the box plot represents the median, box indicates the interquartile range. Black dashed black line represents the WT value. *** denote *P*-value between the two datasets is statistically significant (two-tail *t*-test, *P* < .001). If the difference is not significant, it is represented by n.s. The null hypothesis states that the hypothesised mean difference between the two datasets for which the *P*-value is calculated is zero, implying the RF^CcdB^ values for the two datasets are similar. (D) Frequency distribution of the residue averaged Relative Fitness^CcdB^ scores.

The RF^CcdB^ of the synonymous mutants mainly ranges from 0.5 to 2, a reason to consider mutants having RF^CcdB^<0.5 or RF^CcdB^>2 to be deviated from the WT phenotype. We classified the mutants into three classes i.e., hyperactive, neutral, and inactive based on the variant score, i.e., RF^CcdB^ less than 0.5 is ‘hyperactive’, between 0.5 and 2 is ‘neutral’, and more than 2 is ‘inactive’. A similar cut-off for the hyperactive mutant class was also found using a *k*-means clustering algorithm (Figure S2 C). The entire dataset was also divided into four structural categories (1) CcdA interacting residues, (2) only Gyrase binding residues (excludes the overlapping CcdA interacting residues), (3) buried site residues, and (4) exposed non active-site residues (excludes both CcdA interacting and Gyrase binding site residues) to comprehend functional properties of the mutants in these regions (Figure 1 C). The RF^CcdB^ scores for CcdA interacting mutants (Median = 0.5) were significantly different from all other classes (two-tail t-test, *P* < .001). A significant difference exists between the mean RF^CcdB^ values for the CcdA interacting site mutants and the exposed CcdA non-interacting site mutants (median = 1.5) (two-tail t-test, *P* < .001). However, RF^CcdB^ scores for buried site (median = 3) and only Gyrase binding site mutants (median = 2) lie in a similar range i.e., primarily giving an inactive phenotype. This kind of screening distinguishes CcdA interacting residues from other classes but buried residues cannot be discriminated from Gyrase binding site residues.

To ensure that each mutant amino acid contributes equally to the overall average of the RF^CcdB^ levels for each position, we first averaged the RF^CcdB^ scores for all synonymous mutants of each mutant amino acid and then further averaged over all the mutant amino acids at each position (Figure 1 D). Only CcdA interacting residues have avgRF^CcdB^<0.5, suggesting that mutations at the CcdA binding site result in the severest phenotypes. Mutational analysis indicates 52% (475/920) of the hyperactive mutants belong to the CcdA interacting site, 42% (389/920) to the exposed CcdA noninteracting class, and 6% (56/920) were buried. Among exposed CcdA noninteracting class, 30% (275/920) lie proximal (within 8Å) to CcdA residues, whereas 12% (114/920) lie distal to it. *K*-means clustering, that divided the dataset into 2 clusters also point towards CcdA interacting mutant enrichment in cluster 2 (Figure S2 C).

### *In-vivo* activity and *in-vivo* solubility assays mirror deep sequencing data

Phenotypes of selected mutants inferred from deep sequencing data were validated by individual transformations of the point mutants and plating in both CcdB resistant and sensitive strains (Figure 2 A). Plasmids containing the WT operon and an operon with non-functional toxin were used as controls for inferring the phenotype of the mutant relative to the WT, and for accounting for transformation efficiency differences between the two strains, respectively. The WT construct used in this study has a mutation in the putative SD sequence of CcdA because of a restriction site introduced to facilitate cloning of the *ccdA* coding region of the operon (Figure S3 A and Figure 2 A). The growth in the sensitive strain (Top 10) of the WT construct used in this study was compared with a construct without any mutations in the promoter and identical to that present in F plasmid (WTF’ in Figure S3 B and Figure 2 A). We found that the construct without any mutations in the promoter grew similar to a construct with a stop codon mutation in the toxin gene (Y6_TAA in Figure 2 A) whereas the present WT construct grew more poorly compared to the construct with a non-functional toxin. This indicates that modification of the putative SD sequence of CcdA, likely reduces the expression of CcdA, thereby showing reduced growth and increased toxicity relative to the operon present in F plasmid. The construct exhibiting higher toxicity is used as the WT because it enables to screen both inactive as well as hyperactive mutants (Figure 2 A and S3 A). Deep sequencing results were validated by the growth phenotypes shown by 25 individual CcdB mutants in the sensitive strain, which were in concordance to their inferred RF^CcdB^ scores (Figure 2 A). CcdB mutants displaying a hyperactive phenotype did not grow in the sensitive strain suggesting these mutants show higher toxicity than the WT. P72L which has a RF^CcdB^ score of 0.55 shows no growth in the sensitive strain whereas S47V with a score of 0.6 grows weakly in the sensitive strain. This supports use of RF^CcdB^ cut-off score of 0.5 that was chosen to classify mutants as hyperactive.

**Figure 2:**
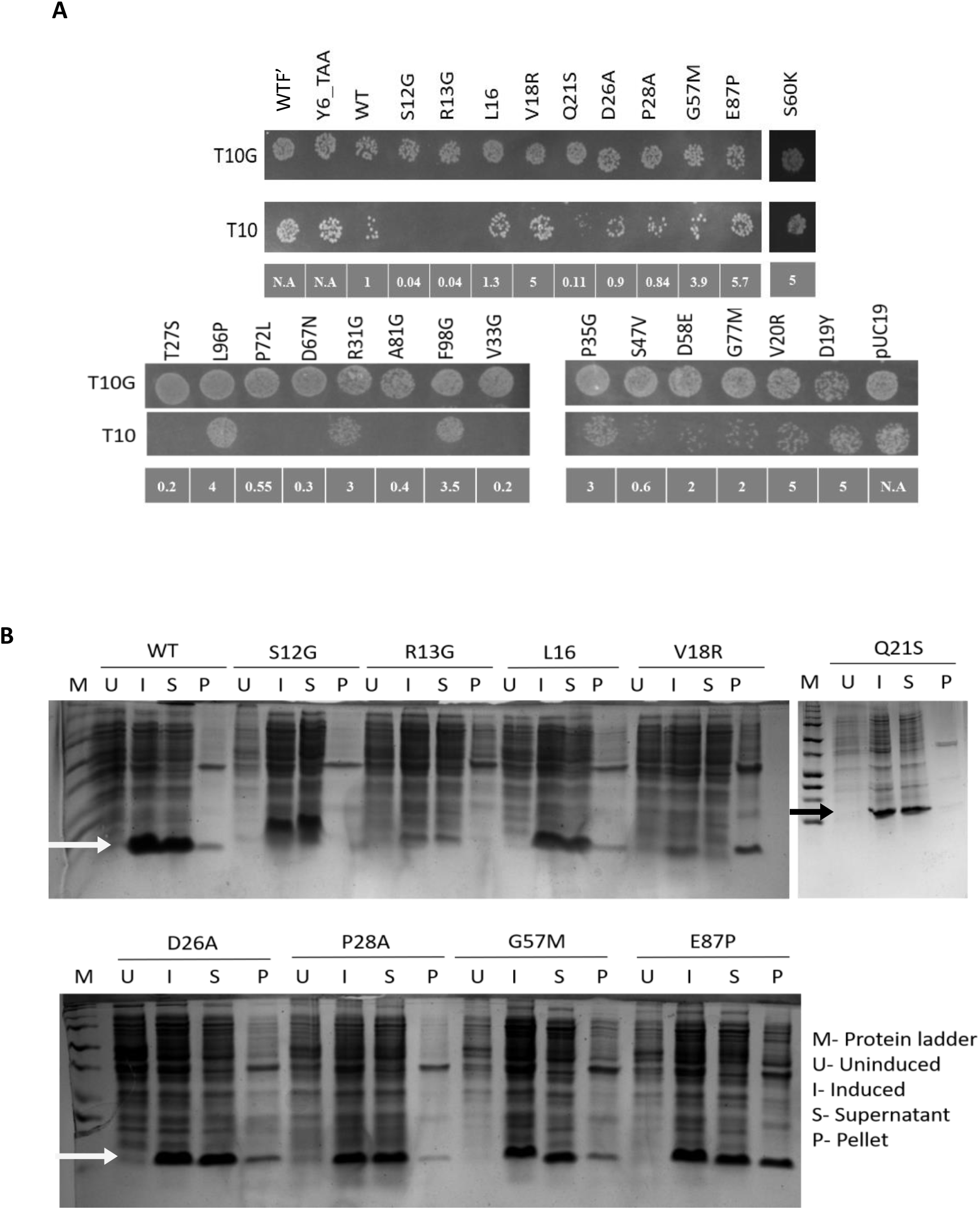
*In vivo* activity and *in vivo* solubility of CcdB mutants validate deep sequencing results. (A) CcdB mutants with different RF^CcdB^ values are selected for individually spotting on LBamp plates. RF^CcdB^ for each mutant spotted has been indicated in the panel below each Top10 panel. N.A is mentioned for controls. (B) *In vivo* solubility is estimated from the relative fractions of protein in supernatant and pellet, determined by densitometric analysis, following 15% SDS-PAGE. The arrow indicates the protein of interest (CcdB). The relative estimates of protein present in the soluble fraction and inclusion bodies for all mutants are shown in Table S3. These experiments were performed in duplicates.

An *in-vivo* solubility assay was conducted for a subset of CcdB mutants (Figure 2 B). In this assay, CcdB was expressed in the absence of CcdA, under control of the pBAD24 promoter^24^. The percentage solubility of CcdA interacting mutants (S12G, R13G, D26A and P28A) was comparable to the WT CcdB protein. L16, a synonymous mutant, also showed *in vivo* solubility comparable to the WT. V18R a buried mutant, likely due to its misfolded nature, mainly was found in inclusion bodies. Although Q21S is a buried mutant, it showed a hyperactive phenotype both in the deep sequencing data as well as in an individual spotting assay, and showed *in vivo* solubility comparable to WT. Percentage solubility of G57M and E87P, that had shown inactive phenotypes in the deep sequencing data, decreased drastically, with 30% and 50% of the induced fractions for G57M and E87P, respectively targeted to inclusion bodies (Figure 2 B). Thus, the *in vivo* solubility assay is consistent with deep sequencing results.

### RF^RelE^ and RF^CcdB^ phenotypic scores provide structural insights

We engineered a construct with the *ccd* promoter upstream of the reporter gene i.e., RelE, a toxin of the RelBE TA operon^25^. We standardised the reporter assay by manipulating the putative Shine Dalgarno sequence to modulate the expression of RelE toxin (see methods in Supplementary Material) (Figure S4 A and B). A significant difference between the growth levels of Top10Gyr co-transformed with WT *ccdAB* operon and consensus RelE, relative to co-transformation with a mutant *ccdAB* operon containing a stop codon CcdB mutant and consensus RelE was observed at 37°C, in both LB media (2-fold difference) and minimal media (5-fold difference) (Figure S4 C). At 42 °C and 45 °C, the reporter was less sensitive possibly because heat shock induced chaperones might help in folding of CcdA, enhancing its binding to the promoter/operator region even in the absence of CcdB, thus decreasing the difference in the growth between the WT and CcdB stop codon mutant. Since the CcdB mutants should not affect the amount of RelB antitoxin produced in the cells, we transformed the CcdB NNK library in the background of the RelE reporter gene in Top10Gyr strain, to avoid any toxic effects resulting directly from the CcdB toxin. In this system, RelE expression and toxicity is regulated by the level of binding of the CcdAB complex to the operator upstream of the RelE reporter. The reporter thus provides a measure of the amount of CcdAB complex within the cell (Figure 3 A). Each CcdB variant was assigned a variant score defined as ‘Relative Fitness^RelE^’ (RF^RelE^) (Eq.7, see methods). A high correlation (r = ∼0.94) was found between the two biological replicates of the CcdB NNK libraries prepared in the RelE reporter strain using a read cut-off of 100 in the resistant strain, indicating that sequencing errors have largely been removed from the analysis (Figure 3 B). The dynamic range of RF^RelE^ scores is much lower than RF^CcdB^ scores and the RF^RelE^ score of WT is 1. Thus, a variant with RF^RelE^<1 was considered to have increased RelE toxicity, suggesting that the variant has a lower amount of CcdAB complex relative to cells expressing WT CcdAB. A variant with a variant score of more than 1 was considered to have similar to higher levels of CcdAB, relative to cells expressing WT CcdAB. Additionally, a similar cut-off was obtained when the entire dataset was divided into two clusters by k-means clustering algorithm (Figure S5).

**Figure 3:**
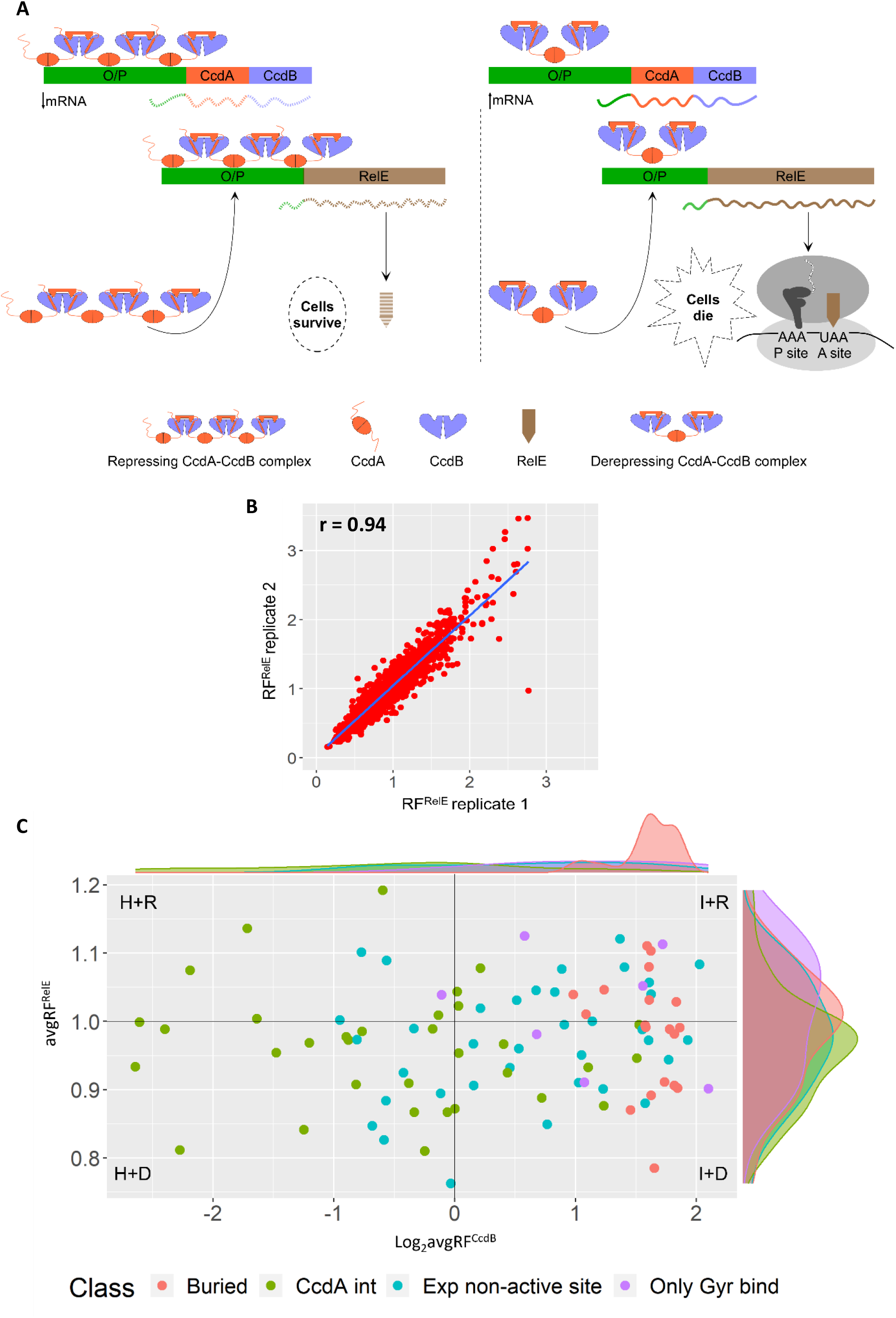
RelE reporter assay to quantitate the relative amounts of CcdA:CcdB complex for different mutants. (A) Schematic of the RelE reporter system. When CcdA is in excess of CcdB (left panel), a repressing complex is formed that binds to the ccd O/P, repressing transcription of the RelE reporter. Decreased mRNA levels and subsequent decrease in RelE is depicted by dashed lines. When CcdA is not in excess of CcdB (right panel), a derepressing complex is formed, resulting in increased transcription of the RelE toxin, causing cell death. Increased mRNA levels and subsequent increase in RelE is depicted by solid lines. (B) Correlation between the two biological replicates for each CcdB mutant for RF^RelE^ values. (C) Corresponding values of Relative Fitness^CcdB^ and Relative Fitness^RelE^ scores averaged over all mutants for each CcdB position belonging to four different classes i.e., Buried site residues (Buried), CcdA interacting site residues (CcdA int), exposed non active-site residues (Exp non active-site) and only Gyrase binding residues (only Gyrase bind). The top horizontal panel is the density spread for different classes based on the avgRF^CcdB^ scores. The right vertical panel depicts the residue averaged density spread for different classes based on the avgRF^RelE^ scores. The black lines represent WT values and divide the graph into four quadrants.

There are 101 residues in CcdB. The first six residues were eliminated because they were used for cloning the previously generated site-saturation mutagenesis library^26^ in the operonic context. Here, we classify the remaining residues based on four structural categories, i.e., 33 CcdA interacting, 7 only Gyrase binding (positions that interact exclusively with GyrA), 19 buried and 36 exposed non active-site residues. The mutants were further classified into four mutational categories based on the RF^CcdB^ and the RF^RelE^ levels defined as (1) Hyperactive and Derepressing (RF^CcdB^<0.5 and RF^RelE^<1), (2) Inactive and Repressing (RF^CcdB^>2 and RF^RelE^>1), (3) Inactive and Derepressing (RF^CcdB^>2 and RF^RelE^<1), and (4) Hyperactive and Repressing (RF^CcdB^<0.5 and RF^RelE^>1) (Table S1). 60% (223/369) of the mutants belonging to the ‘hyperactive and derepressing’ (RF^CcdB^<0.5 and RF^RelE^<1) class fall at the CcdA binding interface. 31% (114/369) of the mutants belonging to this class are exposed CcdA non-interacting mutants, of which 19% (69/369) mutants are proximal to the CcdA chains whereas 12% (45/369) are distal to it. The remaining 7% (26/369) are buried mutants. Enrichment of CcdA interacting mutants and mutants proximal to CcdA chains in this class indicate that these mutants are impaired in binding to CcdA. 30 out of a total of 33 residues at the CcdA binding site are found in this class. Surprisingly, Gyrase binding site mutants (14%) were not predominantly enriched in the ‘inactive and repressing’ class (Table S1). The buried site mutants (36%) dominate over other structural categories in the ‘inactive and derepressing’ class (Table S1), implying that several mutations at buried residues result in misfolding of the protein which in turn hampers its binding to GyrA as well as CcdA. Interestingly, CcdA interacting residues (62%) are also enriched in the ‘hyperactive and repressing’ class (Table S1).

We further averaged all the variant scores of the CcdB mutants for each position obtained from the RelE reporter strain and sensitive strain. (Figure 3 C). Only CcdA interacting residues (7/7) are enriched in the class with avgRF^CcdB^<0.5 and avgRF^RelE^<1 (Figure 3 C), suggesting that the majority of mutations at the CcdA interacting site impede CcdB binding to CcdA, resulting in enhanced amount of CcdB *in vivo*, thereby causing cell death. 14% (2/14) of residues belonging to the class with avgRF^CcdB^>2 and avgRF^RelE^>1 are Gyrase binding site residues. (Figure 3 C). 44% (11/25) of the residues found in the class with avgRF^CcdB^>2 and avgRF^RelE^<1 are buried site residues (Figure 3 C). A large number of buried site mutations were well tolerated in the operonic context (Figure 3 C), likely because CcdA was able to relieve the folding defect of the protein. However, further investigation is required. Therefore, although mutations at both the Gyrase binding site and buried site display an inactive phenotype in Top10, we can still discriminate Gyrase binding site residues from buried residues as Gyrase binding site mutants of CcdB repress RelE expression more efficiently than the buried site mutants, presumably because buried site mutants reduce the amount of properly folded CcdB that can complex with CcdA.

### Structural and mutational data help delineate residue specific contributions to binding

Examination of the phenotypes displayed by CcdB mutants in the sensitive strain revealed that CcdA interacting mutants primarily displayed a hyperactive phenotype (Figure S6 A). Based on the cell growth phenotype analysed for the same class of mutants in the RelE reporter strain, the mutants were classified into two categories, repressing (RF^RelE^>1) and derepressing (RF^RelE^<1) the RelE reporter gene expression. Mutants with RF^RelE^<1 were likely defective in binding to CcdA whereas the mutants with RF^RelE^>1 are associated with larger amounts of the CcdAB complex relative to WT (Figure S6 B). Mutations in 8-14, 23-30, 41-46 and 64-72 residue stretches are enriched in the ‘hyperactive and derepressing’ class. Mutants at residues 12, 13, 28, 30, 42, 43, 46 and 66, CcdA interacting positions are most enriched in this class (>10) (Figure 4 A and B). Residues 24, 25, 26, 96 and 101 are part of both the CcdA and Gyrase binding sites. Amongst these positions, mutants at position 25 and 26 are found in this class whereas most of the mutants at residues 24 and 96 display an inactive phenotype (Figure 4 A). Most mutations at residue 101 are missing in the current dataset (Figure 4 A). The data suggest that I24 and L96 make more critical contacts with Gyrase rather than CcdA. Mutants at residues 24, 87, 88, 91, 92, 95, 96, 99 and 100 of CcdB, which predominantly interact with residues at the Gyrase binding interface do not show much impairment in CcdA binding (Figure 4 A-D). This suggests that residues important in CcdA and Gyrase binding are largely independent from each other.

**Figure 4:**
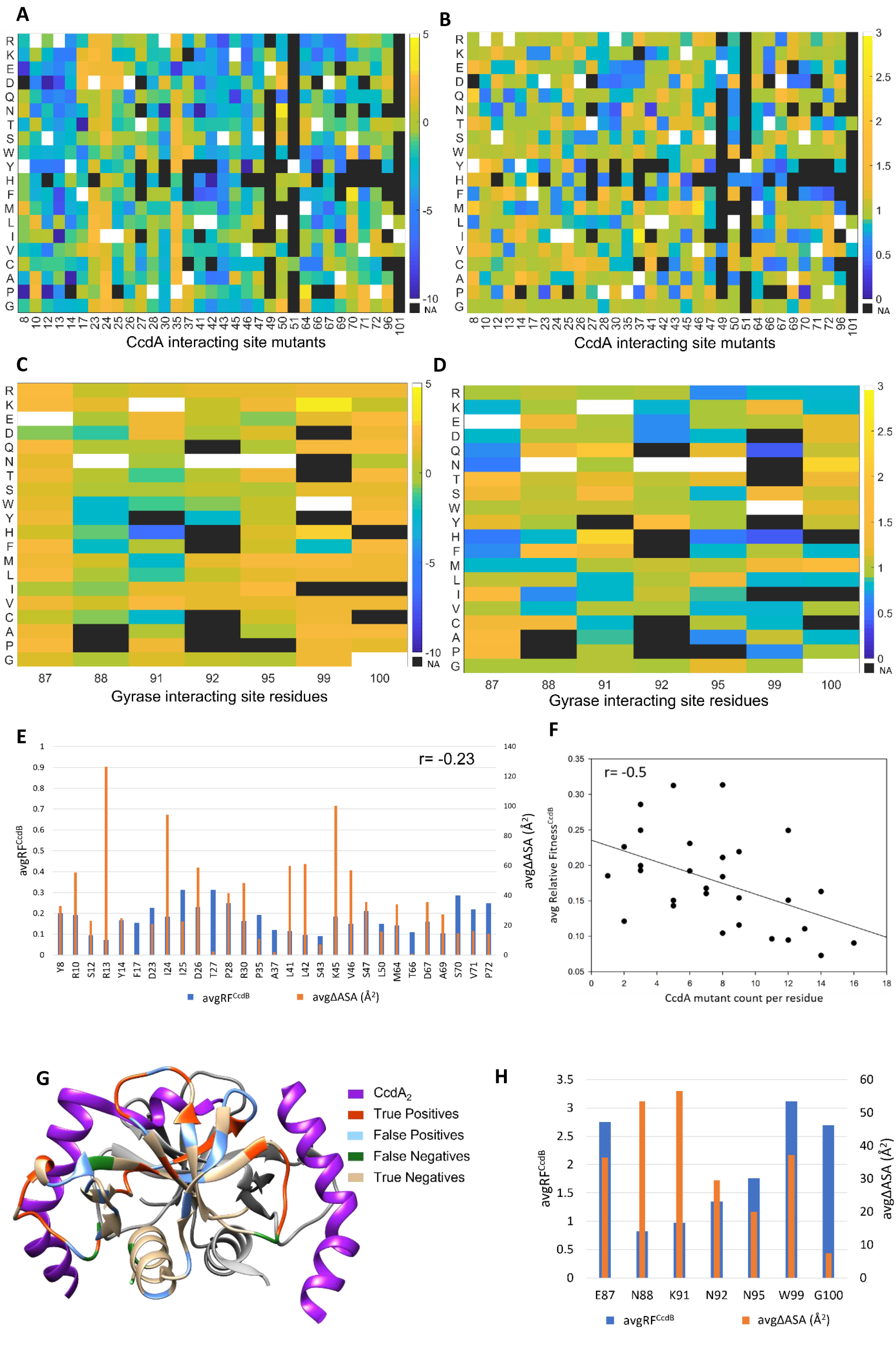
RF^CcdB^ and RF^RelE^ heatmaps and structural correlates. (A-D) Heat maps of Relative Fitness^CcdB^ and Relative Fitness^RelE^ scores. Blue to yellow color gradation represents increasing RF^CcdB^ and RF^RelE^ values. RF^CcdB^ scores are shown in log scale (Log_2_RF^CcdB^) whereas RF^RelE^ scores are shown in linear scale. No data is shown by black colour, i.e., NA (Not Applicable). WT residue at each position is indicated in white. (A) Heat map of RF^CcdB^ scores for CcdA interacting site mutants. (B) Heat map of RF^RelE^ scores for CcdA interacting site mutants. (C) Heat map of RF^CcdB^ scores for Gyrase binding site mutants. (D) Heat map of RF^RelE^ scores for Gyrase binding site mutants. (E) Comparing the avgRF^CcdB^ for each CcdA interacting residue which has RF^CcdB^<0.5 and RF^RelE^<1 with the accessible surface area buried upon complex formation with CcdA. (F) Correlation between avgRF^CcdB^ scores and mutant count enrichment at the CcdA interacting site residues with RF^CcdB^<0.5 and RF^RelE^<1. (G) Residues with RF^CcdB^<0.5, RF^RelE^<1 and ≥3 such mutants/residue are mapped onto the crystal structure of CcdB (PDB ID: 3G7Z)^17^. One monomer of CcdB is shown in light grey while the residues identified as True Positives (orange), False Positives (blue), False Negatives (green), and True Negatives (tan) from the mutational phenotypes are mapped on the other monomer. Majority of the False Positives lie proximal to CcdA chains (<8Å of the CcdA chains). (H) Comparing avgRF^CcdB^ values with ΔASA for residues that bind only to DNA Gyrase.

We averaged the accessible surface area of the CcdB surface buried upon complex formation across the two chains of CcdA and also averaged the RF^CcdB^ scores over mutants for each CcdA interacting residue belonging to the ‘hyperactive and derepressing’ class. Although a weak negative correlation was obtained between the two parameters for each CcdA interacting residue (r=-0.23), we observed that residues that bury more than 30Å^2^ surface area upon CcdA binding have an average RF^CcdB^ score less than 0.2, i.e., ∼8 fold higher toxicity than the WT (Figure 4 E). We also calculated the number of mutants belonging to this class for a given CcdA interacting residue and defined it as mutant count enrichment for a CcdA interacting residue. Average RF^CcdB^ score and the mutant count enrichment for CcdA interacting residues is negatively correlated (∼-0.5), suggesting the residues that have a large number of mutants falling in this class and simultaneously display low average RF^CcdB^ score are the most important residues for CcdA binding (Figure 4 F). These analyses identify S12, R13, F17, R30, L41, L42, S43, V46 and T66 as critical residues for CcdA binding (Table S2). All the residues in the 41-43 stretch interact with E54 of CcdA i.e., part of the 52-55 helical stretch of CcdA, implying E54 in CcdA is likely an important residue for binding to CcdB. CcdA interacting residues get highlighted when mutants belonging to the ‘hyperactive and derepressing’ class with an additional constraint of ≥3 mutants/residue, are mapped onto the 3G7Z crystal structure^17^ (Figure 4 G).

The Relative Fitness scores based on CcdB and RelE toxicity also help to delineate the most important residues for Gyrase binding. Based on the RF^CcdB^ and avgRF^CcdB^ scores, most of the mutants at these residues are inactive while a few are partially active (Figure 4 C and H). As estimated from the RF^RelE^scores, most Gyrase binding site mutations repress RelE expression except for mutations at residue 99 which is buried in free CcdB (Figure 4 D). Gyrase binding induces conformational changes in CcdB^18, 27^. E87 and G100 were identified as the most important residues for Gyrase binding because most mutants at these positions repress RelE expression while showing an inactive phenotype (Figure 4 C-D and H). Surprisingly, N88 and K91 that show significant contact with Gyrase (ΔASA>50Å^2^) have avgRF^CcdB^ scores close to 1, implying that these interactions are not critical for Gyrase binding (Figure 4 H).

### Substitution preferences at functional residues versus buried residues

To further analyse substitution preferences, we grouped amino acids into the following categories aliphatic (A, C, I, L, M, V), aromatic (F, W, Y, H), polar (N, Q, S, T), and charged (D, E, K, R) with G and P into separate categories^2^. Mutations at most residues for the CcdA binding site are found in the ‘hyperactive and derepressing’ class (Figure S7 A). For the Gyrase binding site, mutations to M, S and T are enriched in the ‘inactive and repressing’ class (Figure S7 B). We observed mutations to charged (R, K, E, D), and Glycine residues are primarily enriched for buried site residues belonging to the ‘inactive and derepressing’ category (Figure S7 C). Mutations to hydrophobic residues at buried positions are expectedly not enriched in this class. Therefore, by combining phenotypic scores and careful examination of the mutational pattern, discrimination of Gyrase binding residues from buried residues is possible from mutational data alone.

### Evaluating performance

To avoid bias present at the codon level, we averaged RF^CcdB^ and RF^RelE^ scores over synonymous mutations and then averaged over all mutants found for a given position. Simply examining the distribution of positions belonging to different structural categories into the four mutational categories based on these two averaged RF^CcdB^ and RF^RelE^ scores per residue was not sufficient in discriminating between CcdA interacting, only Gyrase binding and buried residues. We therefore used the absolute values of RF^CcdB^ and RF^RelE^ and impose the additional constraint that a minimum number of mutants (3 or 5) per residue should have the appropriate RF^CcdB^ and RF^RelE^ values. The above criteria (summarised in Table 1) provided relatively accurate assignment of residues to the different structural categories. Results of statistical tests applied to evaluate the performance of this method for predicting CcdA interacting site, Gyrase binding site and buried site residues are mentioned in Table 1. As we enhance the stringency by increasing the number of mutants per residue in the same category (Table 1), the sensitivity decreases whereas the specificity increases.

**Table 1:**
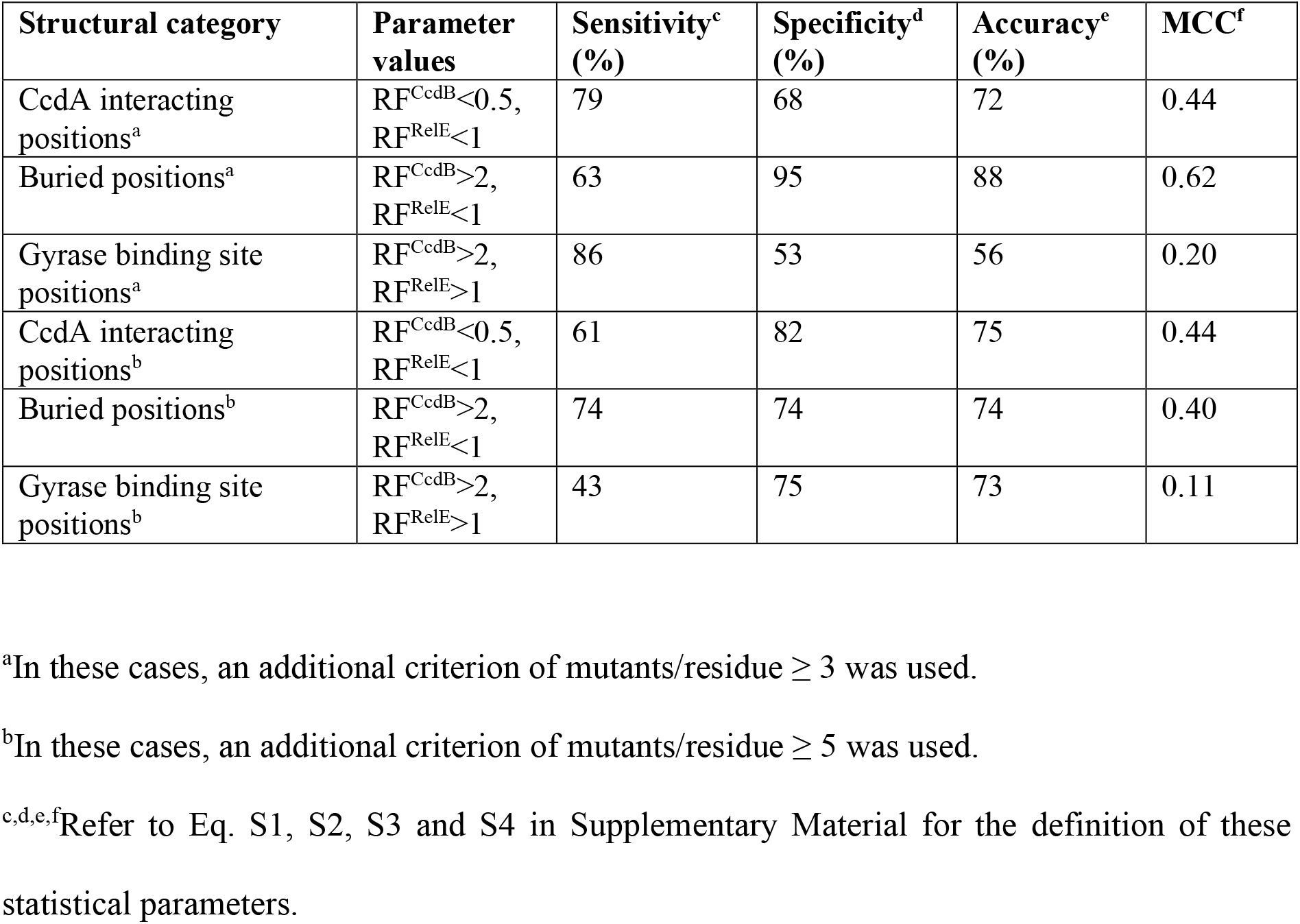
Prediction of active-site and buried positions solely from mutational data, based on RF^CcdB^ and RF^RelE^ scores.

### Characterisation of selected mutants validate their inferred molecular mechanism

A subset of CcdB mutants were individually purified and identities were confirmed by measuring their mass by ESI Mass Spectrometry (Figure S8). Equal concentrations of all the CcdB proteins were used to measure the thermal stability of CcdB variants in the absence and presence of CcdA using nanoDSF (Figure 5 A and B). The thermal shift in T_m_ upon CcdA binding observed for WT is 12 °C. Thermal shifts observed for D26A, G57M and E87P are 15, 14.5 and 14 °C, respectively i.e., slightly higher than the WT. These mutants also show decreased RelE reporter expression *in vivo* (RF^RelE^>1) (Figure 5 C). In contrast, R13G, Q21S and P28A show a thermal shift of 2, 8 and 10 °C, respectively (Figure 5 A and B) i.e., less than the WT and also conferred increased RelE expression relative to WT (RF^RelE^<1) (Figure 5 C). These analyses confirm that mutants that display weaker binding to CcdA relative to WT derepress RelE expression whereas mutants that display stronger binding to CcdA than WT repress RelE expression relative to WT. Out of the three CcdA interacting-site mutants characterised, the largest decrease in binding to CcdA was observed for R13G. Reduced binding of the CcdB variant with CcdA was further confirmed by Microscale Thermophoresis experiments in which binding of the CcdB variant R13G was monitored with respect to CcdB WT protein. Since, CcdB has two interaction sites with CcdA, one low affinity site in the micromolar range^17^ and the other high affinity site in the picomolar range^28^, the CcdB protein was titrated in both the concentration ranges with a fixed concentration of fluorescently labelled CcdA peptide. R13G mutant did not bind to CcdA whereas WT protein bound to lower affinity (µM) binding site of CcdA (Figure S9). Binding affinity of CcdB with CcdA could not be determined for the high affinity site due to extremely tight binding of CcdB with CcdA. Gyrase binding studies were caried out for these mutants using SPR. P28A and Q21S mutants that showed a hyperactive phenotype in the deep sequencing data had higher k_on_ rate than the WT protein. R13G showed an anomalous behaviour by weakly binding to Gyrase even though it is not involved in direct contact with GyrA (Figure 5 D).

**Figure 5:**
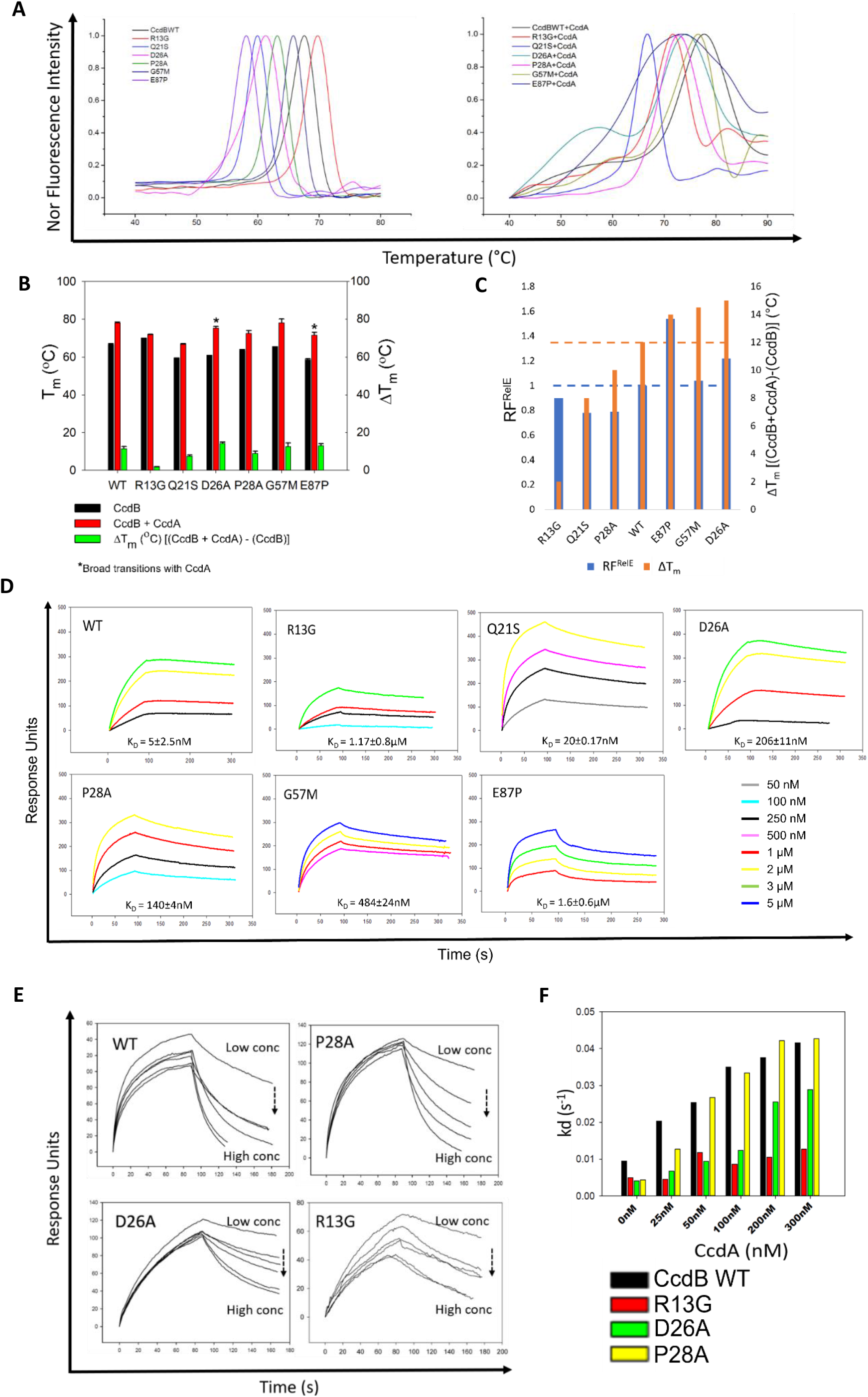
Biophysical characterisation of selected CcdB mutants. (A) Thermal stability of CcdB mutants measured in the absence (left panel) and presence of CcdA (right panel) using nanoDSF. (B) Bar graph (below) represents the T_m_ data. ΔT_m_ is the difference in the T_m_ of the CcdB mutant relative to its complex with CcdA. Error bar signifies standard error between two biological duplicates. (C) Validation of RelE reporter assay. Mutants with RF^RelE^>WT have ΔT_m_>WT while mutants with RF^RelE^<WT have ΔT_m_<WT. (D) Gyrase binding studies of these mutants. For comparative analysis between the mutants, the scale on the y-axis is kept constant. The same concentration for all the mutants is shown by the same colour. The K_D_ values are mentioned below each plot. SPR studies were performed in biological duplicates and, the standard error for the two duplicates is mentioned. (E-F) CcdA_45-72_ mediated rejuvenation of WT and mutant CcdB from their complex with Gyrase. (E) Representative binding sensorgrams of WT and CcdA interacting site mutants. Overlays show the dissociation of WT CcdB and mutants bound to Gyrase with CcdA_45-72_ peptide, concentration increasing from the top to bottom (0 nM, 25 nM, 50 nM, 100 nM, 200 nM, 300 nM). (F) k_d_ (s^-1^) obtained for each mutant as a function of CcdA_45-72_ concentrations. The apparent dissociation rate constants (kd) mediated by CcdA_45-72_ are approximately four-fold lower for the R13G-CcdB-GyraseA14 complex than for WT-CcdB-GyraseA14.

G57M, an exposed CcdA non-interacting mutant was inferred to be inactive from the deep sequencing data. This was validated by SPR studies and several other mutants at the same position also display inactive phenotypes (Figure S6 A). A small to large mutation was not tolerated and hence, the misfolded fraction also increased relative to the WT (Figure 2 B). Phi and psi values of 57^th^ residue for chain A is 53.5 and -128, and for chain B is 78.6 and -106.5, respectively, indicating they lie in the disallowed regions which is only tolerated by Glycine. The 87^th^ position was inferred to be important for Gyrase binding from deep sequencing results. SPR studies show E87P does not bind to Gyrase, further validating the deep sequencing data (Figure 5 D). The characterised data for these mutants is summarised in Table S3.

### CcdA binding residues are also crucial in rejuvenating CcdB from CcdB-Gyrase complex

GyrA14 was immobilized on a CM5 chip and SPR studies were carried out by passing a fixed concentration of CcdB followed by increasing concentrations of CcdA at 25°C. CcdA peptide, residues 45-72 was used for this experiment as it rejuvenates CcdB from the CcdB-Gyrase complex in a fashion similar to full length CcdA protein^18^. When this CcdA peptide was passed over the CcdB-GyrA14 complex, dissociation of CcdB was observed, and the rate of dissociation increased with increasing CcdA concentration (Figure 5 E). SPR studies showed that the dissociation rate of CcdB from GyrA14 in presence of CcdA_45-72_ peptide is approximately 4-fold lower for the R13G variant even at the highest concentration, implying that R13 is a key residue involved in forming a complex with CcdA during or after rejuvenation of CcdB from GyrA. At 25nM CcdA, the dissociation rate of D26A and P28A CcdB mutants is 3 fold and 2 fold lower than WT CcdB, respectively (Figure 5 F). This indicates that WT CcdA dissociates WT CcdB from its complex with Gyrase more efficiently than CcdB with mutations at CcdA binding site residues. Residue 26 interacts with both Gyrase and CcdA. Along with identification of CcdA interacting residues within the CcdB toxin, this strategy can also be used to identify residues important in rejuvenation of CcdB from the CcdB-Gyrase complex.

## DISCUSSION

We have previously employed saturation mutagenesis coupled to deep mutational scanning to infer mutational phenotypes of ∼1600 CcdB mutants, when heterogeneously expressed under the pBAD promoter^1, 2, 26^. In the present study, a comprehensive site-saturation mutagenesis library of the bacterial toxin CcdB, part of a TypeII CcdA-CcdB TA system was generated in its native operon to study mutational effects on organismal fitness. Further, we developed a RelE reporter system that made use of the ability of the CcdB mutants to regulate RelE expression in order to infer the molecular mechanism behind the observed phenotypes. The two phenotypic readouts, one based on CcdB toxicity and the other based on RelE toxicity enabled discrimination between binding site residues to two different ligands, CcdA and DNA Gyrase, as well as between binding site and buried residues.

Mutations can affect activity by either altering the specific activity or total activity, through altering the fraction of the natively folded protein *in vivo,* or by a combination of both^29^. Determining which of the two is the main contributor is a non-trivial task. Computational approaches have been deployed in which sequence-based predicted accessibility scores are combined with mutational sensitivity data to help discriminate between interface residues and buried residues^14^. However, the predictions were only moderately accurate and were limited to predicting contacts for only one interacting partner. In the present study, solely on the basis of the mutational sensitivity displayed by the CcdB mutants and without having to measure the expression levels of individual mutants, we are able to overcome these limitations and predict residues involved in binding to both its cognate partners and discriminate them from buried and exposed noninteracting residues. Mutations of CcdA interacting site residues majorly showed RF^CcdB^<0.5 and RF^RelE^<1 as they were most likely defective in binding to CcdA. Mutational data was in concordance with structural information, and indicated residues 8-14, 41-46 and 62-74 as most important for CcdA binding^17^. CcdB residues in the loop region 8-14 do not contact Gyrase and are important in CcdA binding^17, 18, 30^. Buried site residues were enriched in the class of mutants with RF^CcdB^>2 and RF^RelE^<1 as these mutations often result in misfolding of the protein^2, 31, 32^, thereby hampering their binding to Gyrase as well as CcdA. Most Gyrase binding site residues either showed a neutral phenotype or were enriched in the class of mutants with RF^CcdB^>2 and RF^RelE^>1 as these mutants are well folded^2^ and thus form a complex with CcdA more efficiently than the buried mutants. Exposed non active-site residues are largely insensitive to mutations (0.5<RF^CcdB^<2), a general inference also drawn by other studies^1, 2, 14^. The ones that are mutationally sensitive are either active-site proximal or have RF^CcdB^ values close to the cut-offs. Previous analyses from the laboratory have successfully analysed data from the CcdB mutant library to identify GyrA binding site residues^1, 2^. However, identification of CcdA binding residues was not attempted, as a growth based phenotypic screen had not been developed. In general, utility of mutational scanning datasets increases when combined with appropriate screens to infer molecular mechanisms responsible for the observed phenotypes.

These inferences were further validated via both *in vivo* and *in vitro* studies of the WT and mutant proteins. These studies clearly indicated that a defect in binding to CcdA, resulted in an increase in levels of CcdB in the cell which was sufficient to kill cells more efficiently than the WT. The current approaches that help identify important CcdB residues involved in CcdA binding, will also help identify residues important in rejuvenation of CcdB from CcdB-Gyrase complex. Since, CcdA is an intrinsically disordered protein, identification of these residues can help in understanding the role of intrinsic disorder and allostery in rejuvenation.

Similar approaches can be applied both to other TA systems^33–35^, as well as more generally to other multi-protein systems and should therefore be of general interest. These are especially useful when there are limited sequence homologs available, and where recently developed deep learning approaches^36^ to elucidate protein complex structures do not work well.

## MATERIALS AND METHODS

### Generation of a CcdB site saturation mutagenesis library in its native operon

A previously generated CcdB SSM library^26^ was cloned in pUC57 vector via Gibson Assembly according to the manufacturer’s protocol^37^. The Gibson product was transformed into high efficiency (10^9^ CFU/μg of pUC57 plasmid DNA) electro-competent *E. coli* Top10Gyr cells^38^. The cells were plated on LB agar plates containing 100μg/mL ampicillin, for selection of transformants and incubated for 12 hours at 37°C. Pooled plasmid was purified using a Qiagen plasmid maxiprep kit as per the manufacturer’s instructions.

### Preparation and isolation of barcoded PCR products for multiplexed deep sequencing

The master library was purified from Top10Gyr (resistant strain). The library was then transformed and subjected to selection in both Top10 (sensitive strain) and Top10Gyr harbouring the RelE reporter gene (RelE reporter strain). Pooled, purified plasmid samples from each condition were PCR amplified with primers containing a six-base long Multiplex Identifier (MID) tag. 370 bp long PCR products containing the full *ccdB* gene, were pooled, gel-band purified and sequenced using Illumina Sequencing, on the Hi-seq 2500 platform at Macrogen, Korea.

### Normalization

Read numbers for all mutants at all 101 positions (1-101) in CcdB were analysed. Mutants having less than 100 reads in the resistant strain were not considered for analysis.

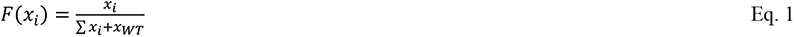

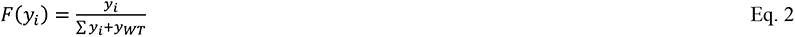

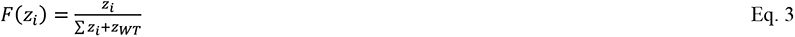

Here, a given mutant is represented by ‘i’ whereas WT is represented by ‘WT’. Number of reads in Top10Gyr resistant strain, Top10 sensitive strain and RelE reporter strain is represented by ‘x’, ‘y’ and ‘z’, respectively. F(x_i_), F(y_i_) and F(z_i_) are the fraction representation of a mutant in resistant, sensitive and RelE reporter strain, respectively.

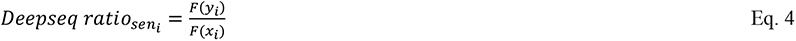

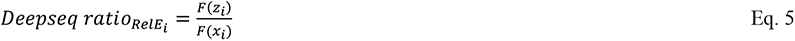

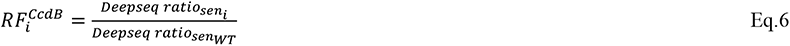

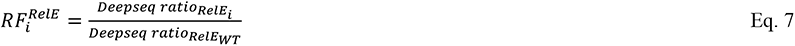

For simplicity, these mutational scores are represented as RF^CcdB^ and RF^RelE^ throughout the text. RF^CcdB^ is based on the CcdB toxicity readout in the Top10 strain while RF^RelE^ is based on the RelE toxicity readout in Top10Gyr strain harbouring the RelE reporter gene. Variant scores of RF^RelE^ were rounded up to 1 decimal place for data analysis. An average of the mutational scores of the two biological replicates for each variant is taken. The two variant scores are generally indicated in linear scale throughout the text.

### *In vivo* activity, expression, and purification of CcdB mutant proteins

Growth assay was carried out for selected CcdB mutants in Top10Gyr versus Top10 *E.coli* strains. Expression and solubility of a subset of CcdB mutants heterologously expressed from pBAD24 vector in Top10Gyr strain was estimated as described previously^2^. These mutants were purified via CcdA affinity chromatography^39^. Protein mass was confirmed using ESI mass spectrometry. All CcdB concentrations reported here are in monomeric units. Purified proteins were further used for nanoDSF, MST and SPR experiments. Detailed description of procedures is mentioned in supplementary material.

## Data Availability

The raw deep sequencing data used in the present study has been deposited in NCBI’s Sequence Read Archive (accession no. SRR17982061).

## SUPPLEMENTARY MATERIAL

Supplementary materials include detailed materials and methods, supplementary tables, and supplementary figures. All supplementary material is available in the file named ‘Supplementary Material’.

## AUTHOR CONTRIBUTIONS

RV and PB designed the experiments. PB performed all the experiments and analysed the data. PB wrote the scripts for data analysis. KM processed the deep sequencing data. RV and PB wrote the manuscript.

## ACKNOWLEDGEMENTS

This work was funded by grants to RV from the Department of Science and Technology, grant number-EMR/2017/004054, DT.15/12/2018) and Department of Biotechnology, grant no. BT/COE/34/SP15219/2015 DT. 20/11/2015, Government of India. We also acknowledge funding for infrastructural support from the following programs of the Government of India: DST FIST, UGC Centre for Advanced study, Ministry of Human Resource Development (MHRD), and the DBT IISc Partnership Program. The funders had no role in study design, data collection and interpretation, or the decision to submit the work for publication. PB acknowledges University Grants Commission, Government of India, for her fellowship. RV is a J. C. Bose Fellow of DST. Nonavinakere Seetharam Srilatha is acknowledged for assisting with the SPR experiments. We thank all members of the RV Lab for their valuable suggestions.

## SUPPLEMENTARY MATERIAL

### Supplementary Materials and Methods

#### Plasmids and Host Strains

The CcdB SSM library was previously generated^1^. Two constructs of the *ccdAB* operon, one with restriction sites and the other without any restriction sites in its native promoter were synthesised in pUC57 vector (3.6kb) at Genscript. The RelE gene downstream of the *ccd* promoter was synthesised in pBT vector at Genscript. Two *E. coli* host strains, i.e., Top10Gyr, strain containing R462C mutation in the GyrA subunit which prevents CcdB from binding to Gyrase^2^ and Top10, strain sensitive to the action of CcdB were used for phenotypic screening of CcdB mutants. The WT and mutant *ccdB* genes were cloned under the control of the P_BAD_ promoter in pBAD24 vector ^3^ and expressed and purified from the Top10Gyr strain.

#### Standardisation of RelE reporter system and generation of CcdB SSM library in a RelE reporter strain

A construct was engineered with the *ccd* promoter upstream of the toxic RelE reporter gene^4^ in pBT vector. Three constructs namely WT RelE, consensus RelE and weak RelE having SD sequence AAGAGG, AGGAGG and AAGTCC, respectively were generated in pBT vector at Genscript. A RelB knockout strain (JW1556 ΔRelB::Kan) from the Keio collection was initially used to standardize the assay. All three constructs were individually co-transformed with the WT *ccd* operon in pUC57 vector in JW1556 ΔRelB::Kan and in Top10Gyr *E.coli* strains. Growth conditions were standardised for the reporter gene by plating cells transformed with the two plasmids at temperatures 30°C, 37°C, 42°C and 45°C. The CcdB SSM library was transformed in Top10Gyr strain resistant to CcdB toxicity harbouring, the RelE reporter gene downstream of the *ccd* promoter with the consensus SD sequence, and plating it at 37°C.

#### Deep sequencing

20% of φX174 viral genomic library was routinely spiked into the pooled sequencing samples. The sequencing was done for two biological replicates. The initial quality of the sequencing data was assessed using FASTQC software. Further analysis was performed using in-house Perl scripts. Sequencing data was analyzed by assigning each read to a particular “bin” based on its MID tag. The downstream primer sequence was used to identify forward and reverse reads in each bin. Only those forward and reverse read pairs which overlap with each other, and together cover the entire *ccdB* gene length, were considered for analysis. The reads were aligned with the wild-type sequence using an in-house script^5^.

#### *In vivo* activity of individual single-site CcdB mutants

Selected single-site CcdB mutants were made in its native operon in pUC57 vector. The amount of plasmid for each mutant was quantitated through nanodrop and confirmed through densitometric analysis, and equal amounts of DNA were transformed in both Top10 and Top10Gyr strain. Different dilutions of the cells were spotted on LB agar plates containing 100 µg/mL Ampicillin and grown at 37°C for 12 hours.

#### Expression and *in vivo* solubility of CcdB mutants

A subset of CcdB mutants were cloned in pBAD24 vector via Gibson assembly^6^. These proteins were heterologously expressed from pBAD24 vector in Top10Gyr strain. Mutants were grown in LB medium, induced with 0.2% arabinose at an OD_600_ of 0.6, and grown for 4.5 hours at 37 °C/180 rpm. Equal numbers of cells (12X10^8^ cells) for all the CcdB variant proteins were taken, sonicated and loaded on 15% Tricine SDS−PAGE gel. Expression and solubility were estimated as described previously^7^.

#### Depth and accessibility calculations

Depth and accessibility of each residue in CcdB was calculated based on the dimeric CcdB structure (PDB ID: 3VUB)^8^, according to the methods described previously^9–11^. A probe size of 1.4 Å was used to calculate accessibility. A residue was defined as buried or exposed if the side chain accessibility is ≤5% or >5%, respectively^12, 13^.

#### Calculation of accessible surface area buried upon complex formation with CcdA and DNA Gyrase (ΔASA)

Accessible surface area values buried upon complex formation with CcdA and DNA Gyrase (ΔASA) were taken from a recent publication^14^. Based on PDB structure 3G7Z, the two chains of CcdA interact with a CcdB dimer. ΔASA is the difference between the solvent accessible surface area of the CcdB residues in free form (3VUB)^8^ and CcdA-bound form (3G7Z)^15^. Similarly, interactions of CcdB residues with GyrA14 were quantitated based on PDB structure 1X75^16^. ΔASA is the difference between the solvent accessible surface area of the CcdB residues in free form (3VUB) and GyrA14-bound form (1X75). We take the average ΔASA value of the two chains for both complexes for comparing with the mutational phenotypes.

#### Evaluation Metrics

We assume the active site residues to represent the positive samples and non active-site residues to represent the negative samples for the prediction of active site residues. On the other hand, for the prediction of the buried sites, we consider the buried residues as the positive samples and the exposed residues as the negative samples. To evaluate the performance of prediction, four evaluation metrics are used in this study: sensitivity, specificity, accuracy, and Matthew’s correlation coefficient (MCC).

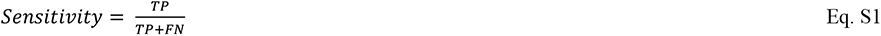

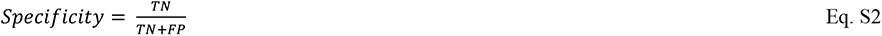

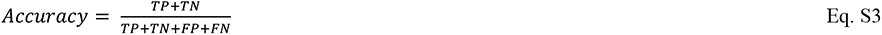

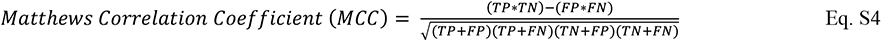

where TP, TN, FP, FN are True Positives, True Negatives, False Positives, and False Negatives, respectively.

#### Statistical analysis

Histograms, k-means clustering, violin, correlation, scatter, and density plots were made using R (version 3.6.3). Heat maps were generated using MATLAB R2019a. Statistical tests and analysis were performed using Microsoft Excel (2016). Values were considered statistically significant for P values below 0.05. P values are provided in appropriate Figures.

#### Purification of CcdB mutant proteins

Proteins were expressed and purified as described previously^17^. 1% of the primary inoculum was inoculated in 500 mL of secondary culture in LB media containing 100 µg/μL ampicillin. Cells were grown to an OD_600_ of 0.6 at 37 °C in LB medium, induced with 0.2% (w/v) arabinose, and grown for 4.5 hours at 37 °C. Cells were harvested by centrifuging at 4000 rpm at 4 °C for 15 min, resuspended in 1/10^th^ volume ice-cold resuspension buffer (10 mM HEPES buffer, 10% glycerol, 1 mM EDTA, and 0.5 mM PMSF, pH 7.4), sonicated on ice, and the total cell lysate was centrifuged at 14000 rpm for 30 min.

CcdA peptide (residues 45-72) synthesised at Genscript, was immobilised on Biorad Affigel-15 as per the manufacturer’s instructions. The immobilization was carried out by incubating beads with the CcdA peptide at pH 7.5, buffered using sodium bicarbonate buffer at 4 °C for 18 hrs, and subsequent blocking by using 1 mM ethanolamine. The protein was purified from the soluble fraction of the lysate by affinity chromatography using immobilized CcdA peptide (residues 45-72). The immobilization was carried out by incubating beads with the CcdA peptide at pH 7.5, buffered using sodium bicarbonate buffer at 4 °C for 18 hrs, and subsequent blocking, using 1mM ethanolamine. The supernatant was loaded onto a CcdA affinity column and incubated at 4 °C for 4 hours, followed by removal of unbound proteins and washing of the column with coupling buffer (MOPS). Protein was eluted with 0.2M glycine, pH 2.5 in 1 mL fractions (a total of 20 fractions were collected) in tubes containing 500 μL of 400 mM Hepes buffer (pH 8.8), to neutralize the protein solution on elution. The eluted fractions were subjected to 15% Tricine SDS−PAGE to check for purity, and the protein was quantitated by comparing the protein band intensity with a standard (Lysozyme).

#### NanoDSF studies

Thermal unfolding experiments of CcdB WT and mutants were carried out by nanoDSF (Prometheus NT.48). The assays were carried out in technical and biological duplicates with 1μM of protein in the absence and presence of 2 µM CcdA_45-72_ peptide in the temperature range of 40 °C–90 °C at 60% LED power and initial discovery scan counts (350 nm) ranging between 5000 and 10,000^18^.

#### Surface Plasmon Resonance (SPR) Experiments

All SPR experiments were performed with a Biacore 3000 (Biacore, Uppsala, Sweden) optical biosensor at 25°C. GyrA14 was used for immobilization at 30 μL/min flow rate for 180s. For Gyrase binding studies, four different concentrations of each CcdB mutant were passed across each sensor surface in a running buffer of PBS (pH 7.4) containing 0.005% Tween surfactant. Protein concentrations ranged from 50 nM to 5 µM. Both binding and dissociation were measured at a flow rate of 30 μL/min. In all cases, the sensor surface was regenerated between binding reactions by one or two washes with 4 M MgCl_2_ for 30 s at 30 μL/min.

For the rejuvenation assay, 250 nM of the WT CcdB and CcdB variant proteins (R13G, D26A and P28A) were then run across each sensor surface in 1X PBS (pH 7.4) containing 0.005% Tween surfactant. For these experiments, the SPR was done in co-inject mode, where the association was allowed for 100 secs, followed by immediate dissociation with different concentrations of the CcdA_45-72_ peptide. Dissociation of CcdB mutants bound to Gyrase was examined by co-injecting increasing concentration of CcdA_45-72_ peptide (0 nM, 25 nM, 50 nM, 100 nM, 200 nM, 300 nM). Rest process is same as mentioned above. The kinetic parameters were obtained by fitting the data to a simple 1:1 Langmuir interaction model by using BIA EVALUATION 3.1 software^14^.

#### Microscale Thermophoresis (MST) studies

The CcdA_45-72_ peptide was fluorescently labelled using the MO-L008 Monolith™ His-tag labelling kit following the manufacturer’s protocol, and diluted in PBST buffer (PBS, pH 7.4, 0.05% Tween-20) to attain a final concentration in the range of 20 – 40 nM to be used in the binding studies. The non-fluorescent CcdB proteins were titrated in a 1:1 serial dilutions manner against the target, CcdA_45-72_ peptide. The normalised fluorescence FNorm is plotted as parts per thousand as a function of CcdB concentration. The dissociation constants (K_D_) were determined employing standard data analysis with MO. Affinity Analysis Software^19^.

---

**Figure S1:**
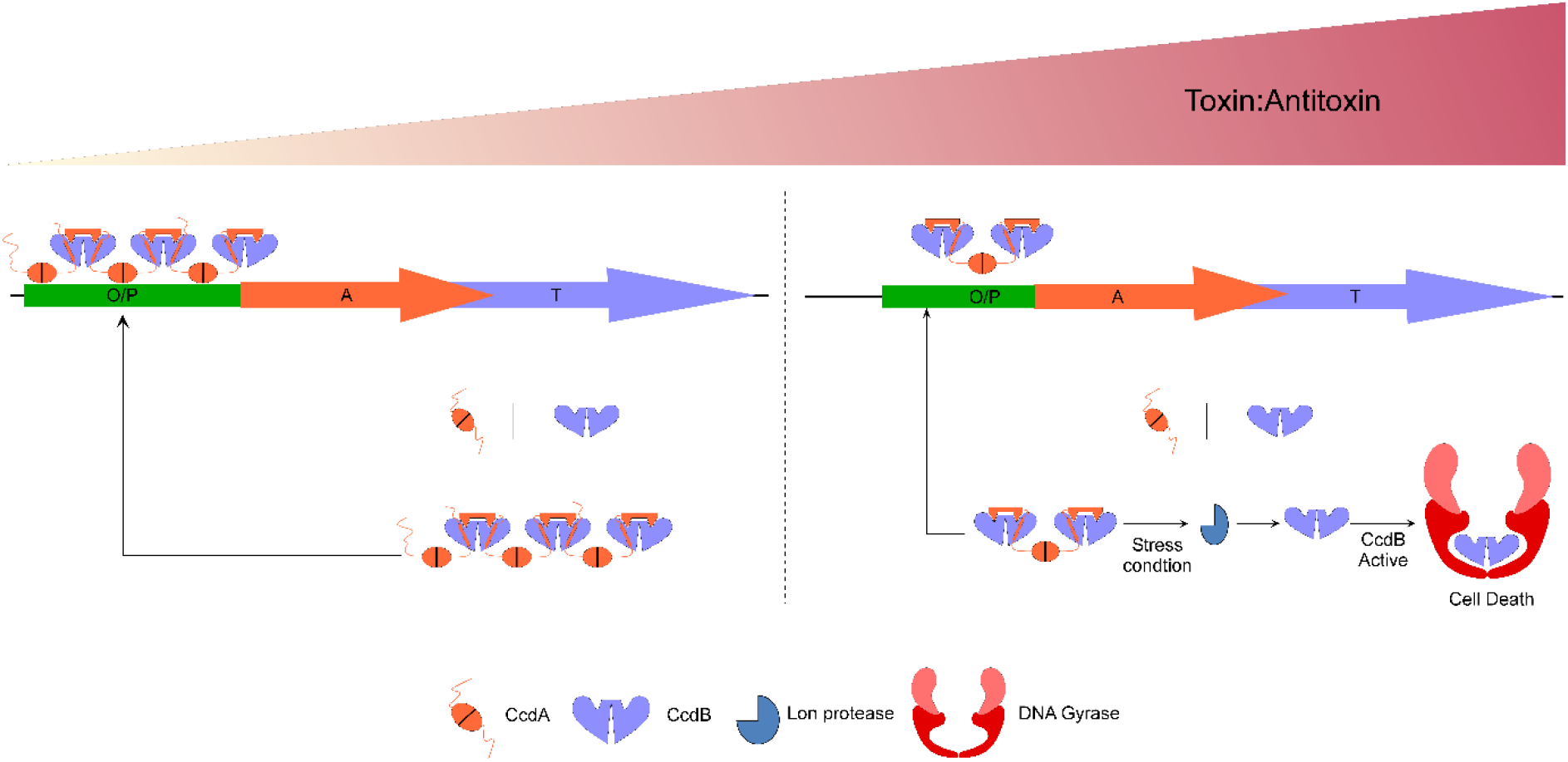
Schematic representation of regulation of CcdAB Toxin-Antitoxin system. Autoregulation is a universal feature of Toxin-Antitoxin systems that determines the toxin:antitoxin ratio within cells. The left panel shows that when [CcdA] > [CcdB], the CcdA-CcdB proteins form an extended multimeric complex that represses transcription of the operon. The right panel shows that when [CcdA] < [CcdB], the CcdA-CcdB proteins form a CcdB:(CcdA)_2_:CcdB heterotetramer that derepresses transcription of the operon.

**Figure S2:**
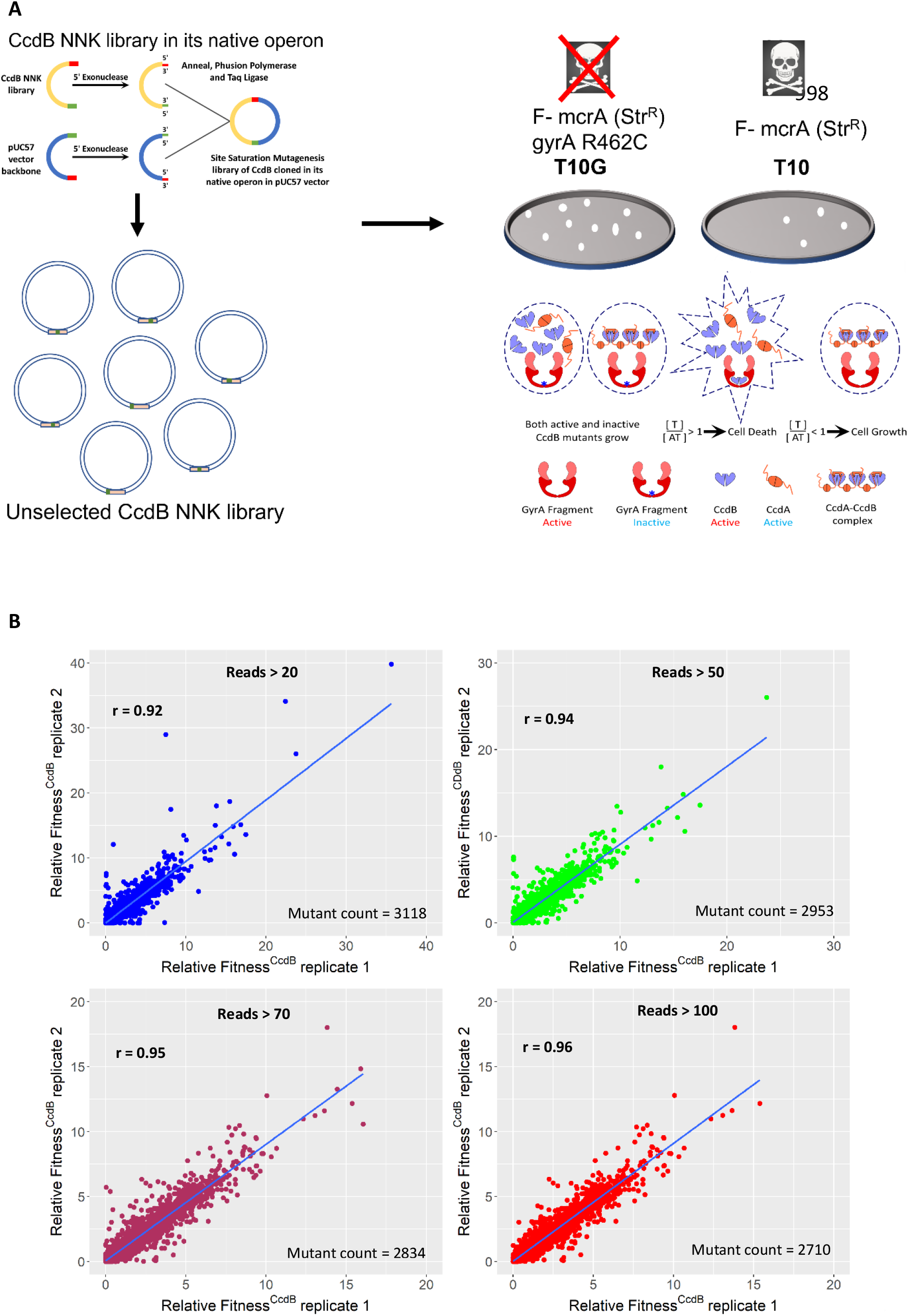
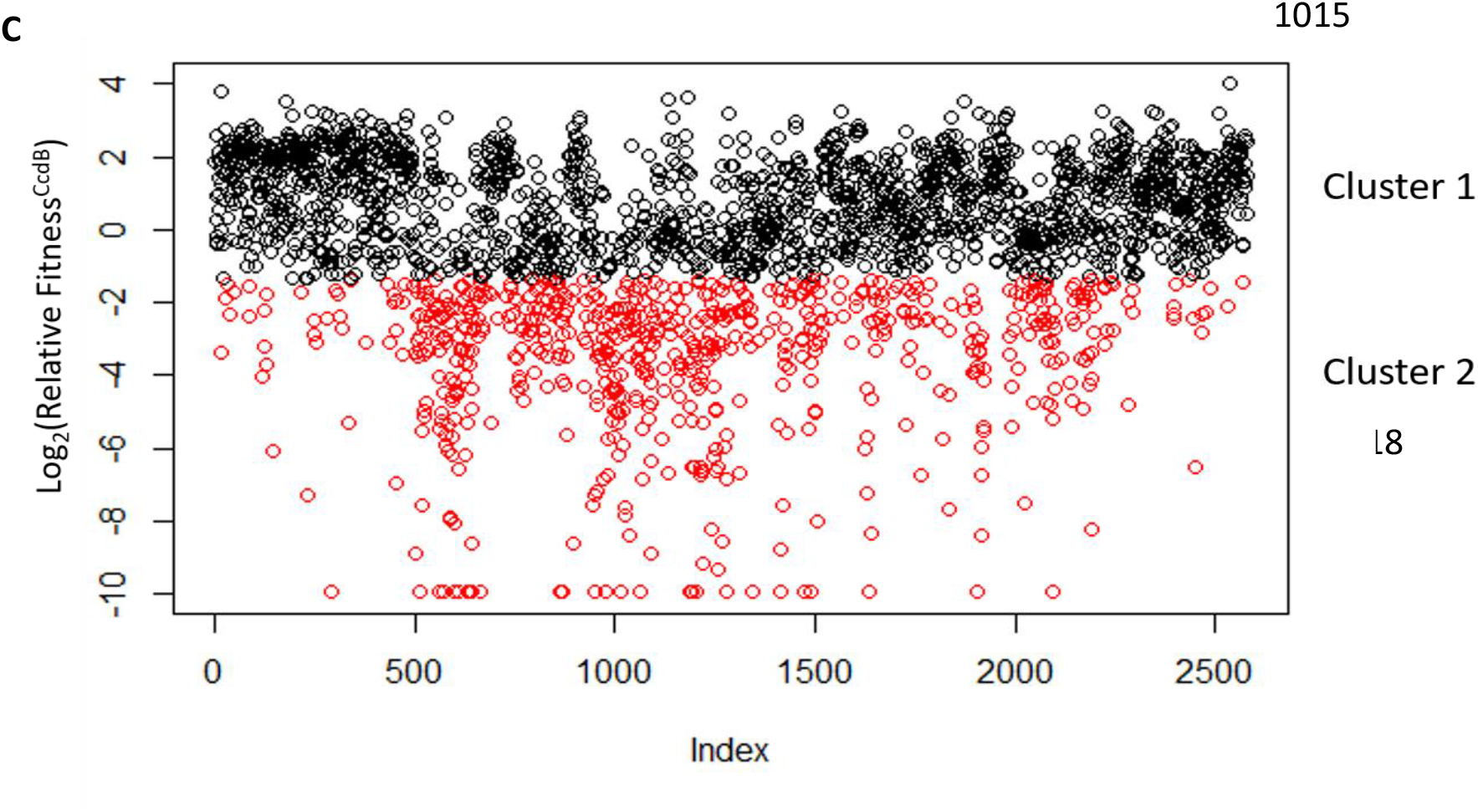
Methodology, reproducibility, and frequency distribution of RF^CcdB^ scores. (A) Pipeline for mutational analysis of CcdB SSM library in native operonic context. All mutants survive in Top10Gyr, CcdB resistant strain whereas only those mutants that bind CcdA and/or fail to bind DNA Gyrase survive in Top10, a CcdB sensitive strain. (B) Correlation between RF^CcdB^ values for the two biological replicates for mutants with different read cut-offs in the resistant strain Top10Gyr. A higher RF^CcdB^ value is associated with lower toxicity and activity of the corresponding mutant. (C) Division of the entire dataset into two clusters based on k-means clustering algorithm. Points falling in cluster 1 are coloured in black whereas points falling in cluster 2 are coloured in red. The boundary separating the two clusters has Log_2_(Relative Fitness^CcdB^) score close to -1. Cluster 1 consists of 462 CcdA interacting mutants, 455 buried mutants, and 964 CcdA non-interacting mutants. Cluster 2 consists of 429 CcdA interacting mutants, 37 buried mutants and, 232 CcdA non-interacting mutants.

**Figure S3:**
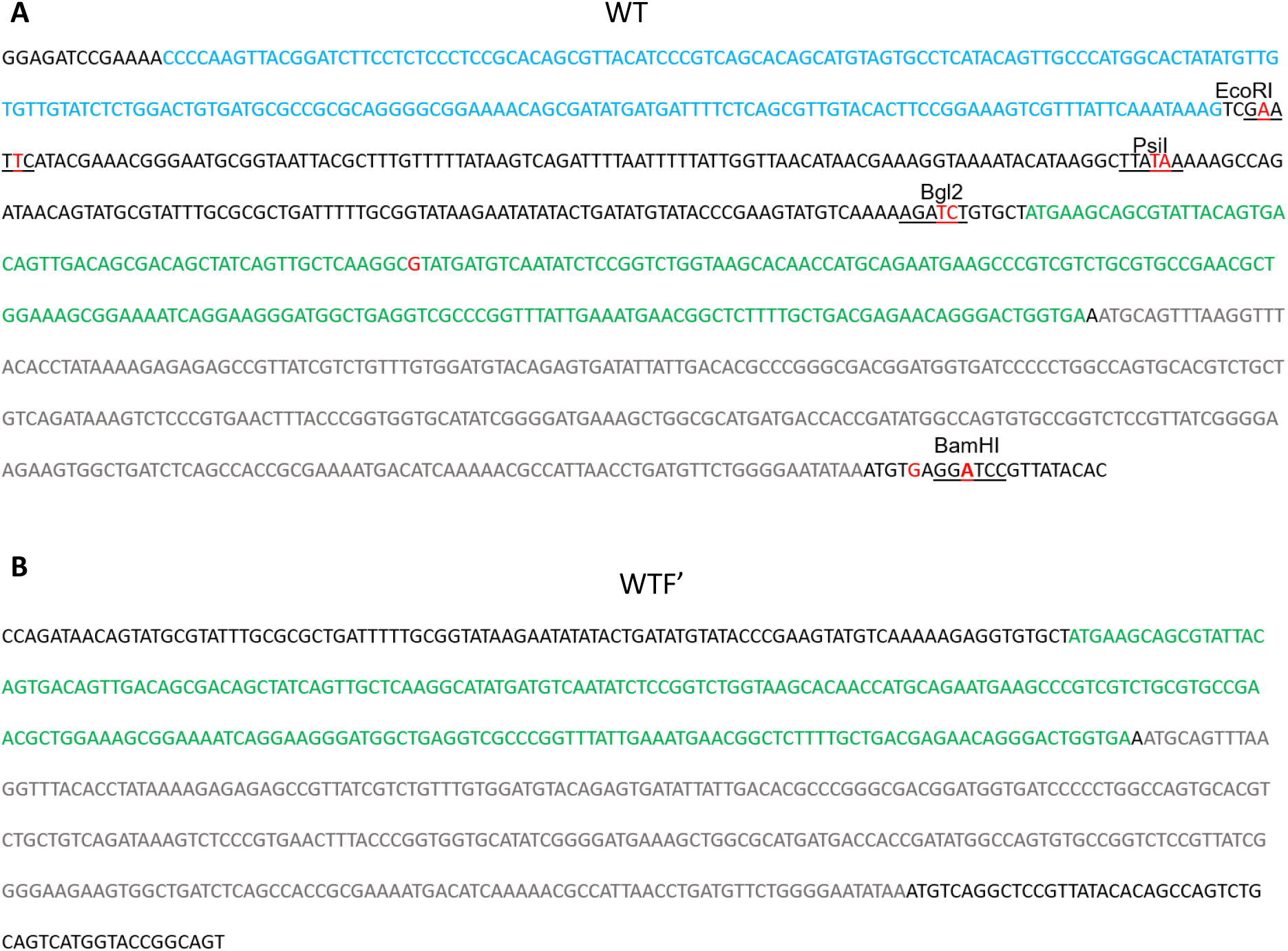
Gene sequence of *ccd* operon used in this study. (A) Sequence of *ccd* operon (968bp) synthesized and cloned at the EcoRV site in pUC57 vector resulting in pUCccd plasmid (F-plasmid coordinates are 45920-46886). Restriction sites have been underlined and marked. *ccdA* coding region is in green and *ccdB* is in grey. Mutations to facilitate cloning are marked in red. Following the *ccdB* gene, the first few residues of the resD gene have been included with a stop at the second codon and mutation to introduce a BamH1 site. This construct is designated WT in this study. (B) Sequence of *ccd* operon (670 bp) without any restriction sites. *ccdA* coding region is in green and *ccdB* is in grey, the first few residues of the *resD* gene have been included to mark the transcription termination site. This is the WT sequence of the *ccdAB* operon found in the F plasmid, we refer to it as WTF’ in this study.

**Figure S4:**
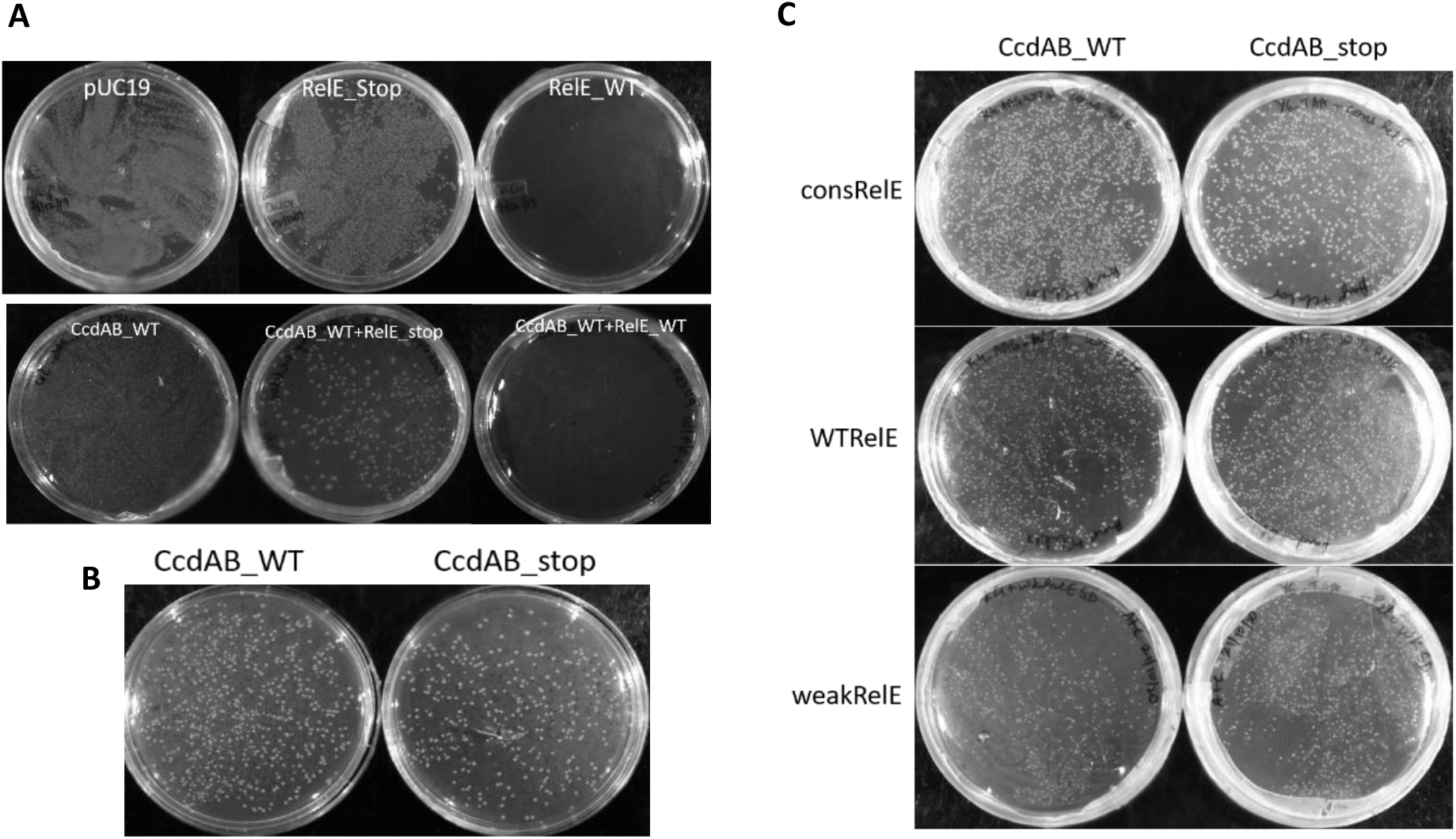
Optimisation of the RelE reporter assay. (A) The pBT plasmid expressing the RelE reporter gene under control of the *ccd* promoter was transformed in JW1556 ΔRelB::Kan. pUC19 and RelE_stop cloned in pBT vector were negative controls, and pBT_RelE_WT was the positive control. CcdAB_WT operon grew equivalent to RelE_stop and pUC19 construct in JW1556 ΔRelB::Kan strain. CcdAB_WT and RelE_stop was co-transformed in this strain. A growth defect is observed when CcdAB_WT and RelE_WT are co-transformed in this strain. (B) Equal growth of CcdAB_WT and CcdAB_stop constructs in Top10Gyr shows that both plasmids are in identical concentration. (C) Growth of Top10Gyr strain when these two constructs are co-transformed with consensus RelE, WT RelE and weak RelE constructs. The maximum difference in CFU was observed when CcdAB_WT and CcdAB_stop were co-transformed with consensus RelE construct, both in LB media as well as minimal media at 37°C.

**Figure S5:**
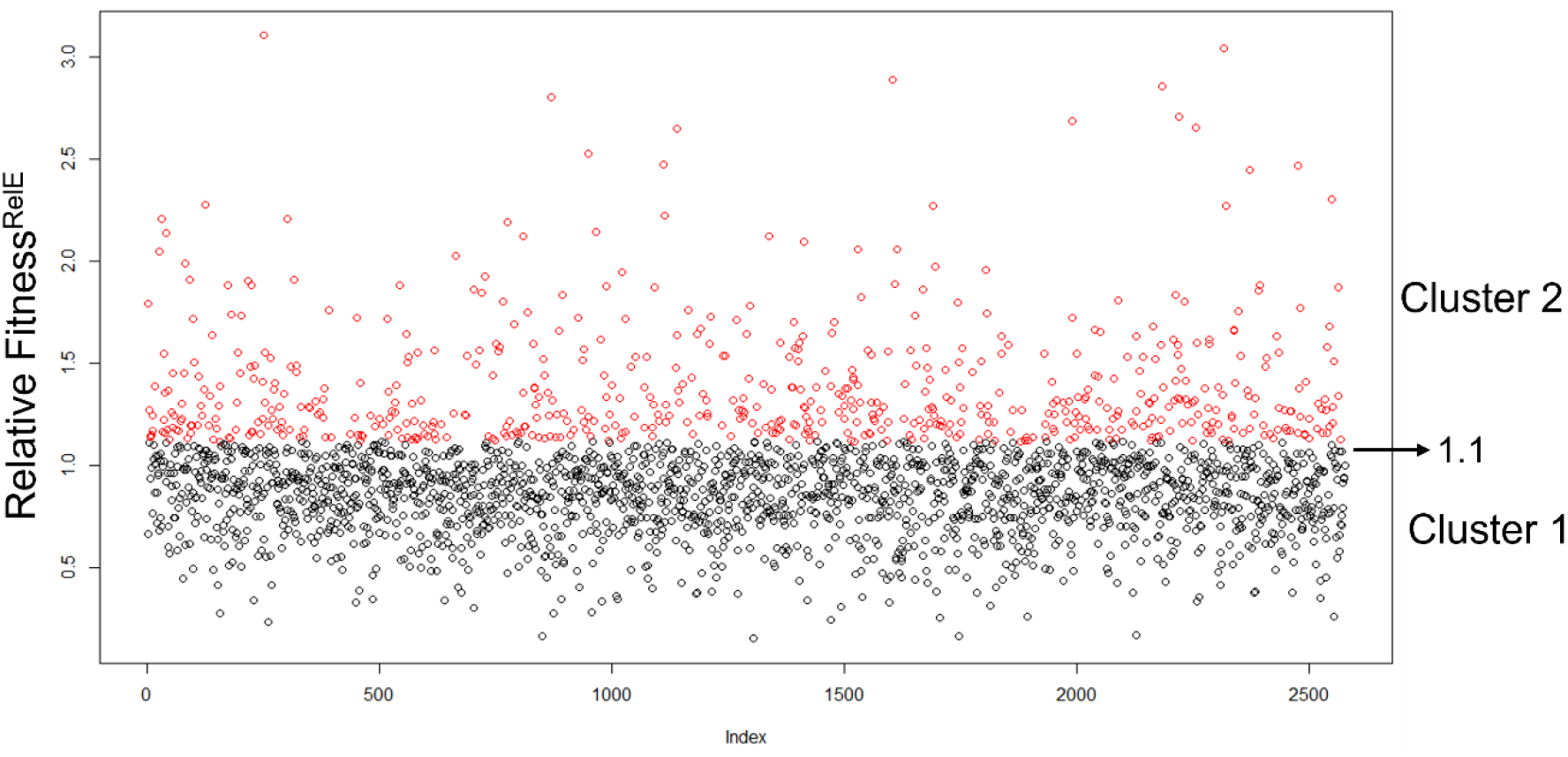
Read cut-off for RelE reporter assay determined by k-means clustering algorithm. Division of the entire dataset into two clusters based on k-means clustering algorithm. Points falling in cluster 1 are coloured in black whereas points falling in cluster 2 are coloured in red. The boundary separating the two clusters has Relative Fitness^RelE^ score close to 1.1. Cluster 1 consists of 684 CcdA interacting mutants, 140 Gyrase binding site mutants, 369 buried mutants, and 741 exposed CcdA and Gyrase non-interacting mutants. Cluster 2 consists of 207 CcdA interacting mutants, 57 Gyrase binding site mutants, 114 buried mutants and, 235 exposed CcdA and Gyrase non-interacting mutants.

**Figure S6:**
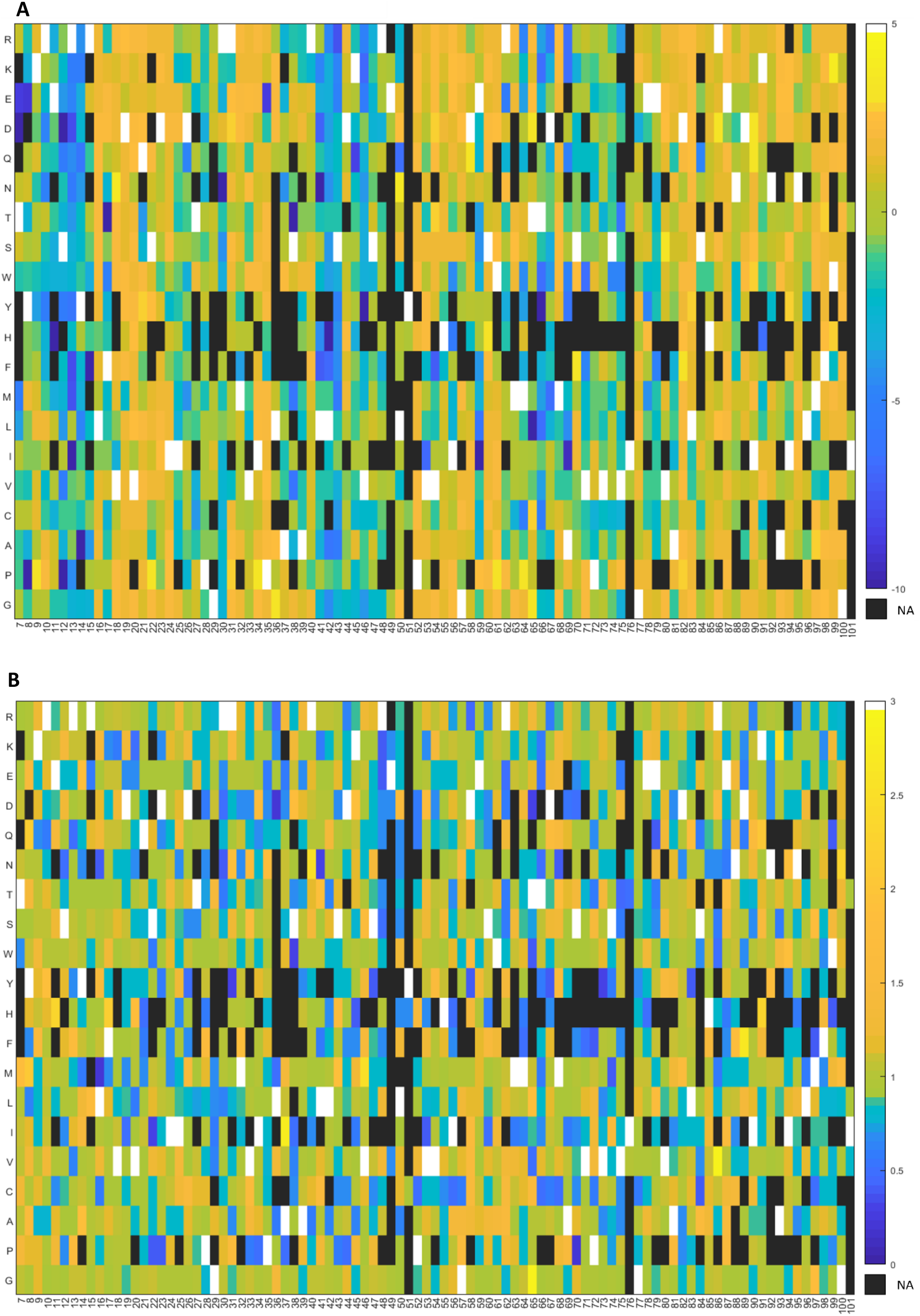
RF^CcdB^ and RF^RelE^ scores for all mutants analysed at the amino acid level. (A-B) Synonymous mutations are averaged for each mutant and then Relative Fitness scores are plotted in the form of heat maps. Residue numbers and mutant identities are shown on x and y axes, respectively. Blue to yellow color gradation represents increasing RF^CcdB^ and RF^RelE^ values. RF^CcdB^ scores are shown in log scale (Log_2_RF^CcdB^) whereas RF^RelE^ scores are shown in linear scale. NA (not applicable) represented by black colour indicates that the corresponding mutant was not observed in the library. WT residue at each position is indicated in white. (A) Heat map of RF^CcdB^ scores for all mutants. (B) Heat map of RF^RelE^ scores for all mutants.

**Figure S7:**
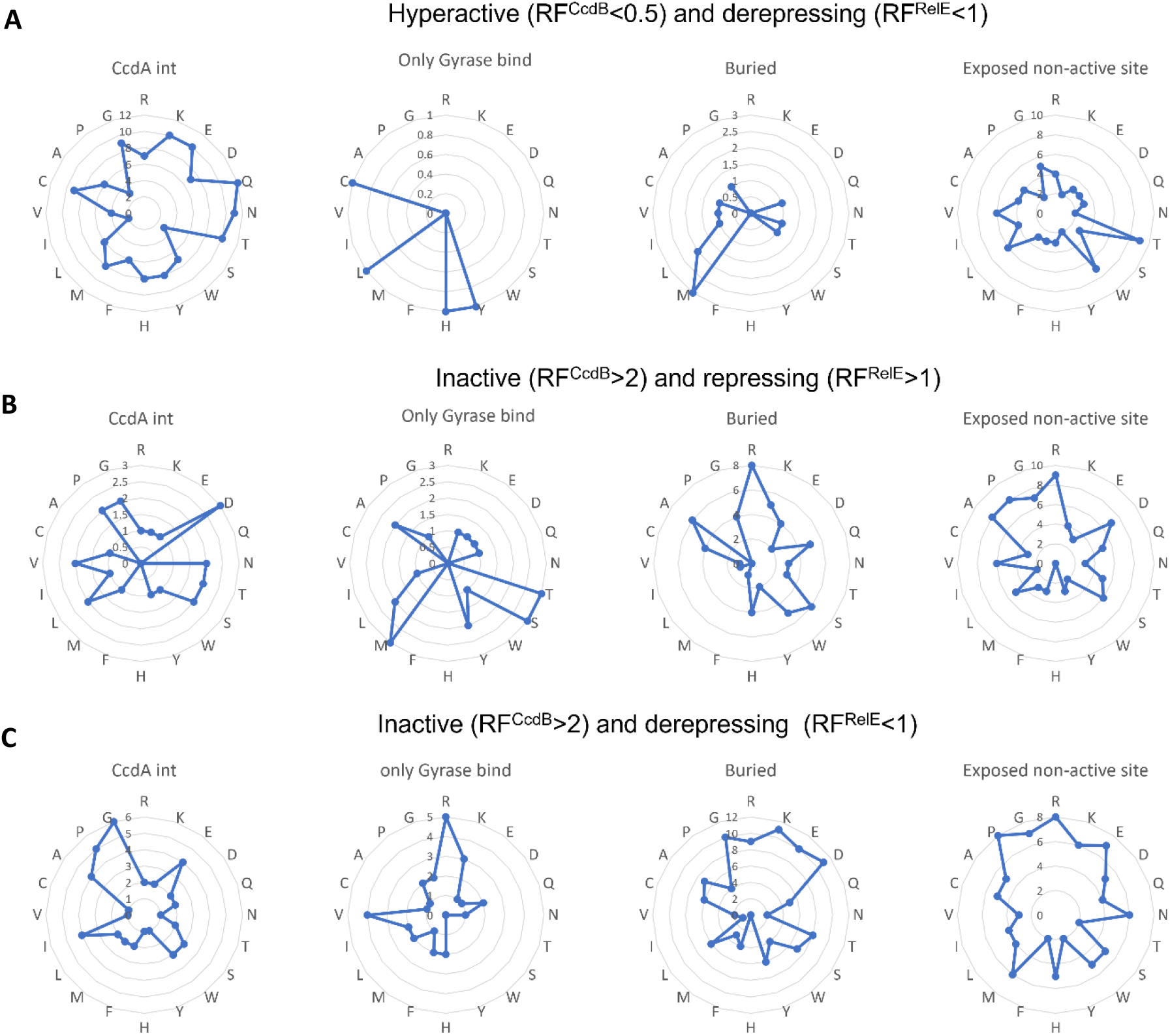
Mutational patterns observed for active site, buried, and exposed non active-site residues. (A-C) Mutant amino acid identity is shown at the circumference of the radar plot. Each circle represents the number of mutants (shown on the vertical axis) obtained for each mutant amino acid for each structural category. (A) Frequencies of mutants with RF^CcdB^<0.5 and RF^RelE^<1. (B) Frequencies of mutants with RF^CcdB^<2 and RF^RelE^>1. (C) Frequencies of mutants with RF^CcdB^<2 and RF^RelE^<1.

**Figure S8:**
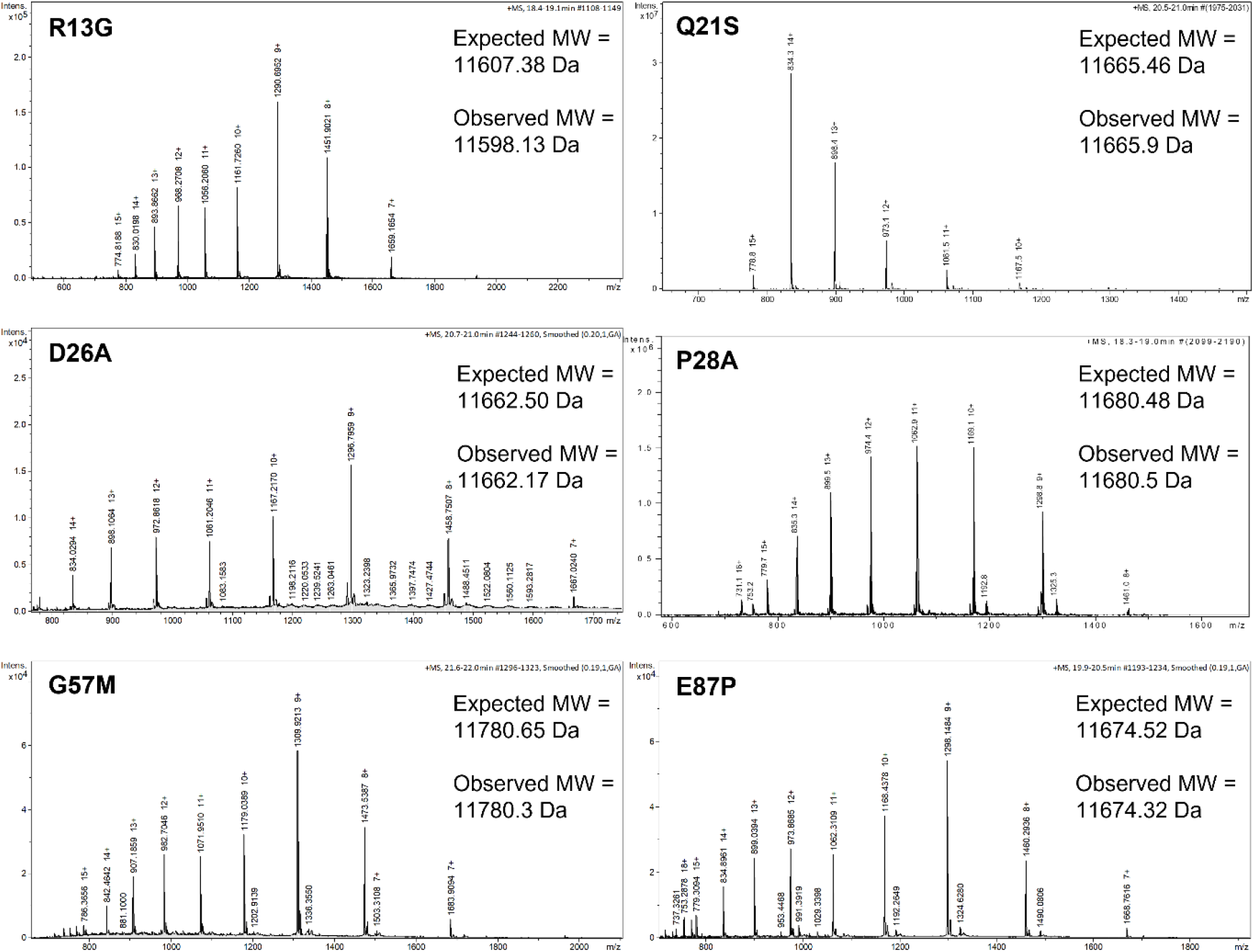
Molecular mass of CcdB proteins confirmed using Mass Spectrometry. CcdB mutant protein peaks are deconvoluted using ESI Mass Spectrometry. Expected and observed molecular weights (MW) are mentioned inside the box.

**Figure S9:**
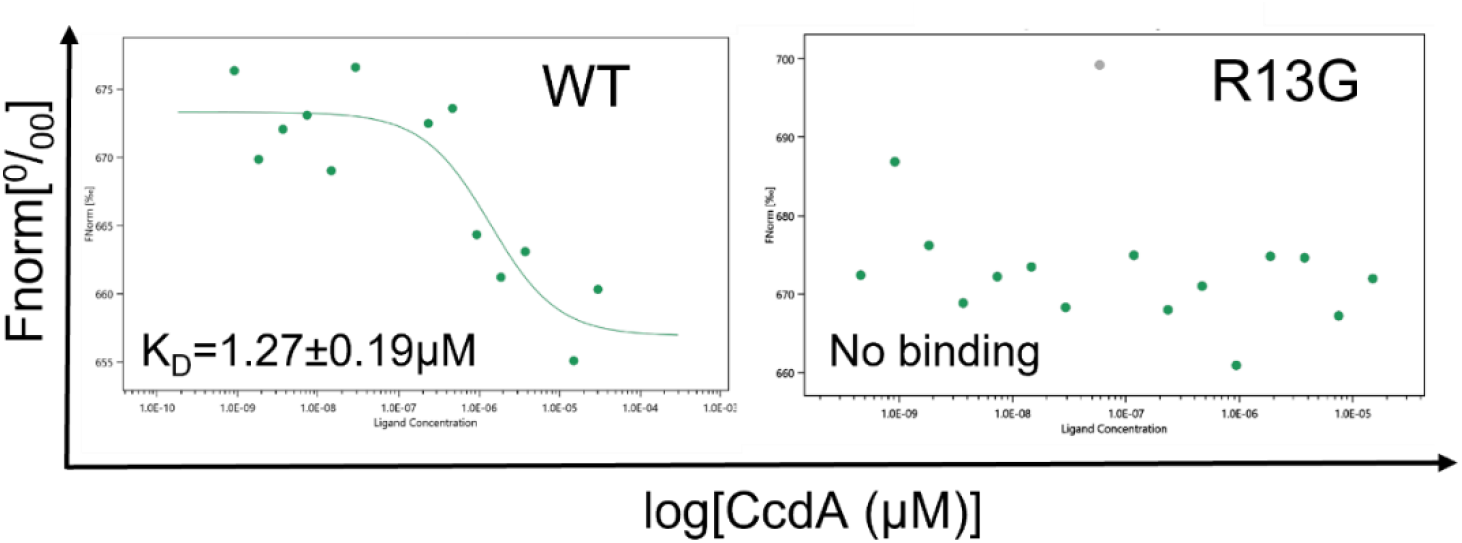
Binding studies of CcdB proteins with CcdA using MST. Binding of the CcdA interacting CcdB variant, R13G to CcdA measured through Microscale Thermophoresis. The experiment was carried out in biological duplicates. Binding of CcdA_45-72_ to the low affinity site on WT CcdB has a K_D_ of 1.27±0.19 µM. Binding of CcdA to the high affinity site on CcdB has a K_D_ in picomolar range which could not be accurately determined. The R13G mutant of CcdB failed to bind CcdA.

**Table S1:**
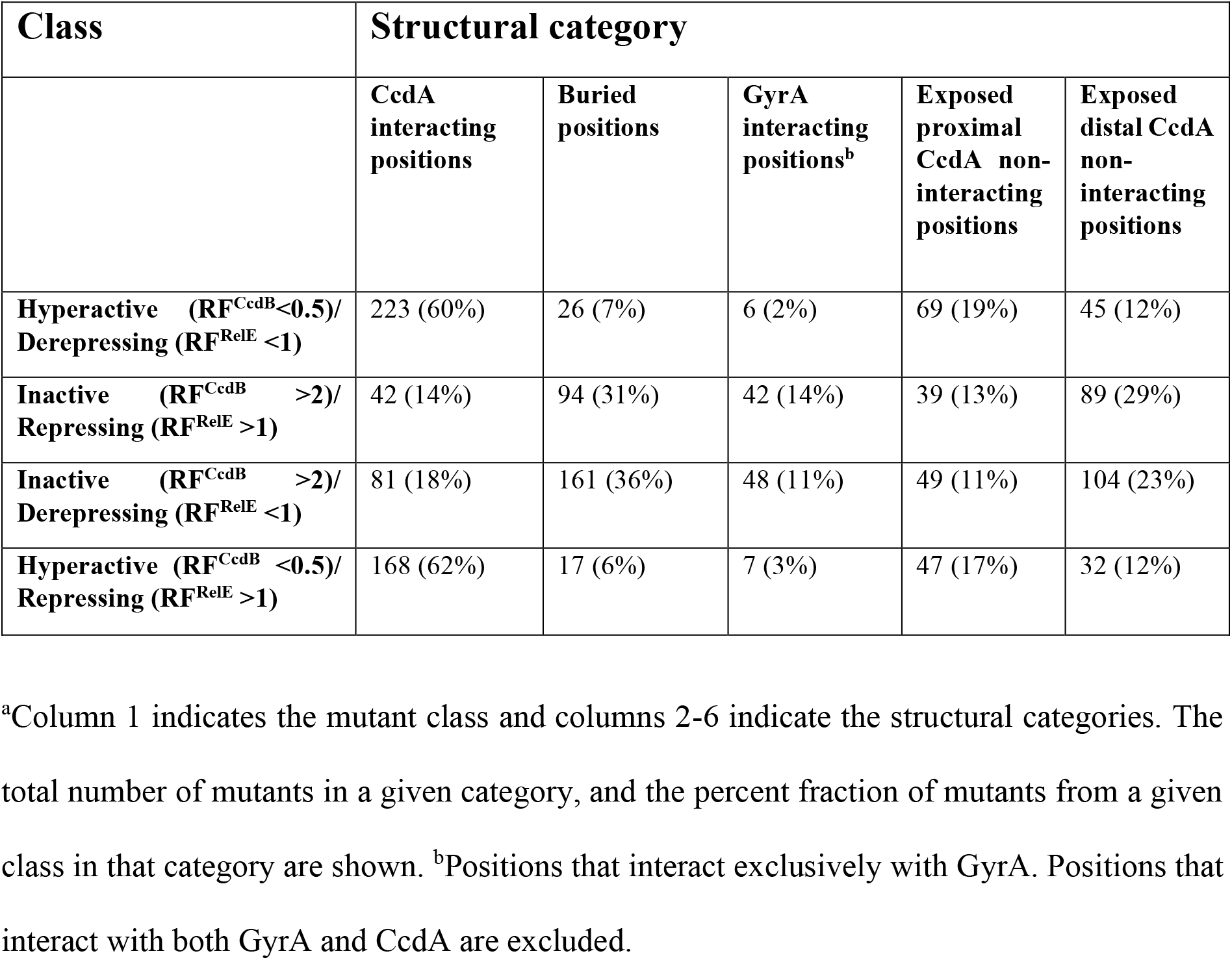
Distribution of CcdB mutants amongst structural categories as a function of CcdB and RelE phenotypic readouts^a^.

**Table S2:**
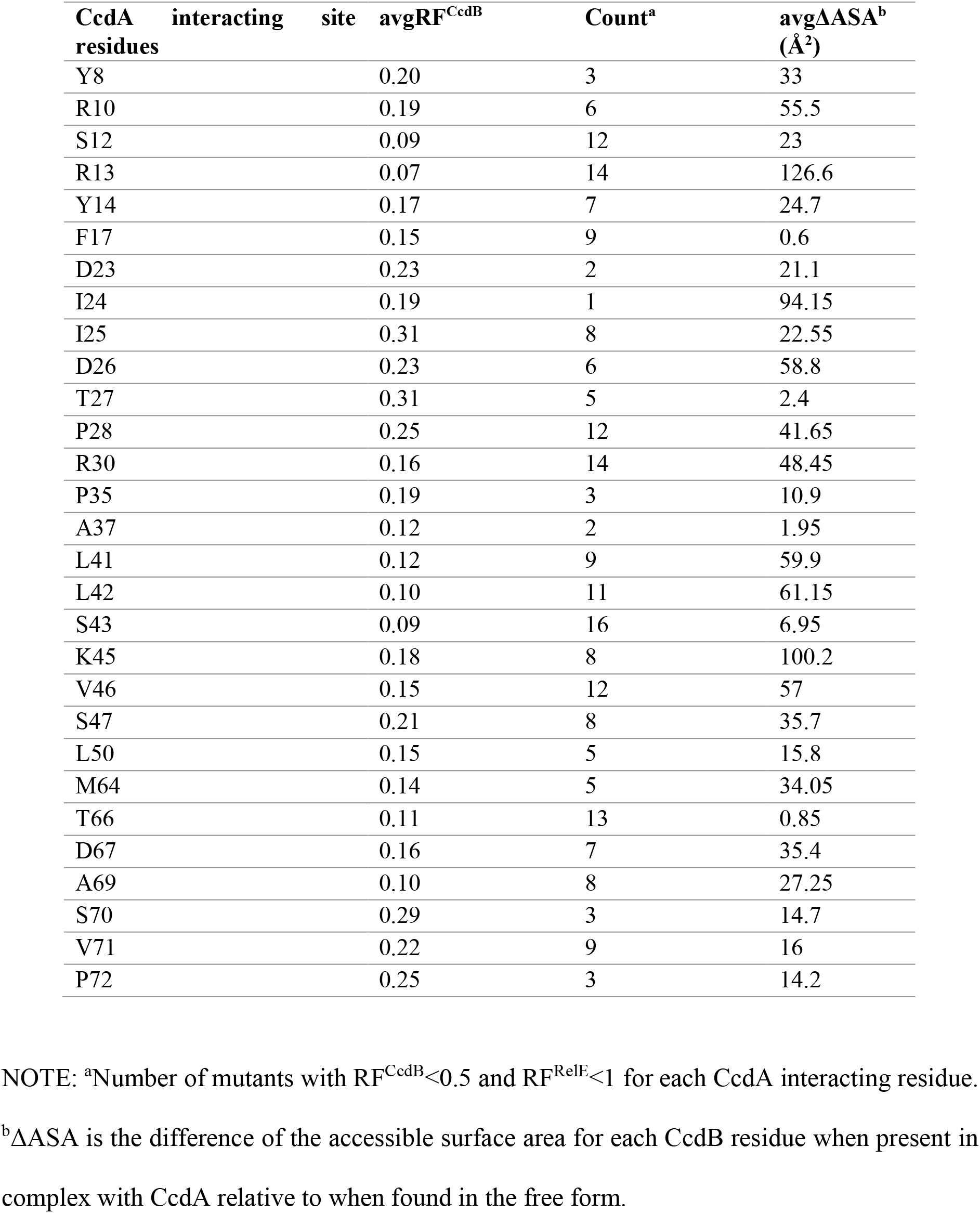
Parameter values of CcdA interacting residues belonging to ‘hyperactive and derepressing’ class.

| **CcdA interacting site** | **avgRF<sup>CcdB</sup>** | **Count<sup>a</sup>** | **avgΔASA<sup>b</sup> (Å<sup>2</sup>)** |
| --- | --- | --- | --- |
| Y8 | 0.20 | 3 | 33 |
| R10 | 0.19 | 6 | 55.5 |
| S12 | 0.09 | 12 | 23 |
| R13 | 0.07 | 14 | 126.6 |
| Y14 | 0.17 | 7 | 24.7 |
| F17 | 0.15 | 9 | 0.6 |
| D23 | 0.23 | 2 | 21.1 |
| I24 | 0.19 | 1 | 94.15 |
| I25 | 0.31 | 8 | 22.55 |
| D26 | 0.23 | 6 | 58.8 |
| T27 | 0.31 | 5 | 2.4 |
| P28 | 0.25 | 12 | 41.65 |
| R30 | 0.16 | 14 | 48.45 |
| P35 | 0.19 | 3 | 10.9 |
| A37 | 0.12 | 2 | 1.95 |
| L41 | 0.12 | 9 | 59.9 |
| L42 | 0.10 | 11 | 61.15 |
| S43 | 0.09 | 16 | 6.95 |
| K45 | 0.18 | 8 | 100.2 |
| V46 | 0.15 | 12 | 57 |
| S47 | 0.21 | 8 | 35.7 |
| L50 | 0.15 | 5 | 15.8 |
| M64 | 0.14 | 5 | 34.05 |
| T66 | 0.11 | 13 | 0.85 |
| D67 | 0.16 | 7 | 35.4 |
| A69 | 0.10 | 8 | 27.25 |
| S70 | 0.29 | 3 | 14.7 |
| V71 | 0.22 | 9 | 16 |
| P72 | 0.25 | 3 | 14.2 |

NOTE: <sup>a</sup>Number of mutants with RF<sup>CcdB</sup><0.5 and RF<sup>RelE</sup><1 for each CcdA interacting residue.
<sup>b</sup>ΔASA is the difference of the accessible surface area for each CcdB residue when present in complex with CcdA relative to when found in the free form.

**Table S3:** Biophysical characterisation of selected mutants from different classes.

| Mutation | RF<sup>RelE</sup> | RF<sup>CcdB</sup> | Solubility (%) | K<sub>d</sub> (GyrA) (nM) | T<sub>m</sub> (-CcdA) (°C) | T<sub>m</sub> (+CcdA) (°C) | ΔT<sub>m</sub> (°C) | Class<sup>a</sup> | Structural category |
| --- | --- | --- | --- | --- | --- | --- | --- | --- | --- |
| WT | 1 | 1 | 90 | 5.5 | 66 | 78 | 12 |  |  |
| R13G | 0.90 | 0.04 | 100 | 1.1 | 70 | 72 | 2 | H+D | CcdA int |
| V18R<sup>b</sup> | 0.97 | 5.03 | 0 |  |  |  |  | I+D | Buried |
| Q21S | 0.78 | 0.11 | 90 | 20 | 58 | 67 | 8 | I+D | Buried |
| D26A | 1.22 | 0.92 | 70 | 206 | 61 | 76 | 15 | H+R | CcdA and Gyrase int |
| P28A | 0.79 | 0.84 | 90 | 140 | 62.5 | 74 | 10 | H+R | CcdA int |
| G57M | 1.04 | 3.99 | 80 | 484 | 65.5 | 78.5 | 14.5 | I+R | Exposed non active site int |
| E87P | 1.54 | 5.75 | 50 | 1.6 | 56 | 73 | 14 | I+R | Gyrase int |

<sup>a</sup>H, hyperactive; I, Inactive; D, Derepressing; R, repressing; int, interacting
<sup>b</sup>V18R is a buried mutant so could not be purified.

